# Adhesion-driven tissue rigidification triggers epithelial cell polarity

**DOI:** 10.1101/2025.03.18.644006

**Authors:** Laura Rustarazo-Calvo, Cristina Pallares-Cartes, Adrián Aguirre-Tamaral, Elisa Floris, Maximilian Hingerl, Camilla Autorino, Arif Ul Maula Khan, Bernat Corominas-Murtra, Nicoletta I. Petridou

## Abstract

The active regulation of tissue material properties via phase transitions is central in morphogenesis. Transitions abruptly occur at critical points in diverse control parameters, including cell density, shape or adhesion. Whether these parameters are interdependent, performing redundant or distinct functions, is unknown. Here we show that co-regulation of multiple control parameters impacts not only tissue deformability, but also cell polarization. We theoretically define a new phase diagram capturing the material states of zebrafish pluripotent tissues and show that they cross simultaneously critical points in cell density, connectivity and adhesion strength. Combining optogenetics, biophysical measurements and quantitative morphometrics, we independently modulate each parameter, identifying adhesion as the main determinant of tissue rheology. Unexpectedly, uncoupling adhesion-driven from density-driven rigidification in amorphous tissues triggers epithelial organization via tricellular junction formation, followed by luminogenesis and apicobasal polarization. Altogether, this work reveals the non-linear dynamics of emergent tissue mechanics as instructive mechanisms of tissue organization.

## MAIN

The functionality of living tissues emerges from the intricate interplay between mechanical and biochemical processes, studied at the intersection of statistical physics, material sciences, and biology. The active regulation of tissue material properties has been recently identified as a key factor establishing spatial and temporal deformation patterns during tissue spreading and folding, body axis elongation, wound healing and metastasis ^1–9^. Recent work has shown that tissue material properties spontaneously and abruptly change between discrete material regimes, e.g., solid-like vs. fluid-like, irrespective of force application ^10–12^. The emergence of collective tissue phases —and their transitions— resemble phases of inert materials, where changes in the material state non-linearly occur when a control parameter describing the interactions of the microscopic constituents reaches a critical point ^13–15^. In contrast to inert materials, living materials display a variety of cellular properties as control parameters of tissue phase transitions, including cell shape, density, connectivity and contact dynamics ^16^. The multiplicity of parameters determining the tissue material state raises the question of which parameters are primarily relevant.

The existence of multiple microscopic cellular mechanisms controlling the macroscopic material state has been observed theoretically and experimentally. For example, cell shape, tension fluctuations and motility ^4,17,18^ and both cell density and connectivity ^18,19^ were shown to trigger material phase transitions in confluent and non-confluent tissues, respectively. The above studies propose a repertoire of simple cell parameters to diagnose the tissue material state, which are regulated by a handful of molecular players, such as actomyosin contractility and adhesion molecules ^20^. The co-existence of multiple control parameters suggests that on the one hand, they might be highly interdependent, but on the other hand, slight differential adaptations of their values can trigger novel types of phase transitions presumably linked to specific biological functions. This raises several questions: (i) Which control parameters are interdependent or redundant? (ii) Can they be uncoupled? and (iii) Does each type of phase transition serve additional biological functions besides defining tissue deformability, depending on the control parameters used ^21^?

We tackle these fundamental questions in the zebrafish early embryo, whose pluripotent blastoderm has been previously shown to undergo a rigidity phase transition, required for its spreading ^5,19^. We theoretically derive a new critical point in contact surface tensions, which we empirically validate within the developing embryo, in combination with previously established critical points in cell density and connectivity ^19,22,23^. We experimentally tune the multiple control parameters independently and disentangle their contributions to both tissue material state and organization. We show that contact surface tension is the major driver of tissue solidification, and, when uncoupled from changes in density, it triggers epithelial features in pluripotent tissues, both at a morphological and molecular level. We therefore show that material phase transitions not only regulate tissue deformation, but also impact the morphogenetic potential of early pluripotent embryonic tissues.

### Multiple control parameters are coupled during an *in vivo* material phase transition

To gain initial insight into which cell parameters regulate tissue material phase transitions *in vivo*, we quantified several cell behaviors in the zebrafish pluripotent embryo (**Fig. 1I**). We have previously measured tissue viscosity in the zebrafish central blastoderm using micropipette aspiration and showed that at pluripotent stages (high, sphere) it has high viscosity, at the onset of morphogenesis (dome) it abruptly decreases its viscosity, and by the time gastrulation is about to start (50% epiboly, shield stage) blastoderm viscosity increases again ^5,24^ (**Fig. 1K**). Using the framework of rigidity percolation theory, we further identified that the above tissue fluidization can be interpreted as a rigid-to-floppy phase transition occurring at a critical point in cell connectivity ^19^. This framework links the material state of a network to a structural element, the size of the Giant Rigid Cluster (GRC), which is the largest subgraph in a network made of viscoelastic links that cannot be deformed without an energy cost (**Fig. S1A, A’**) ^2,25–27^. When connectivity <*k*> reaches the critical value of Maxwellian rigidity <*k*_*c*_>, the GRC abruptly increases from a vanishing size at the subcritical phase (dome stage), to spanning almost all the network at the supercritical phase (high, sphere, 50% and shield stages) (**Fig. S1A, A’**, dashed line, **Fig. S1B, C, Fig. 1J**). Previous theoretical work in hard and soft spheres showed that connectivity can change as a function of both packing fraction and the attractive/repulsive nature of the interactions triggering jamming transitions ^18,23,28^. Assuming that the biological analogues are cell density and cell-cell adhesion strength, we explore how density-dependent vs. adhesion-dependent phase transitions contribute to connectivity changes and the tissue material state.

**Figure 1:**
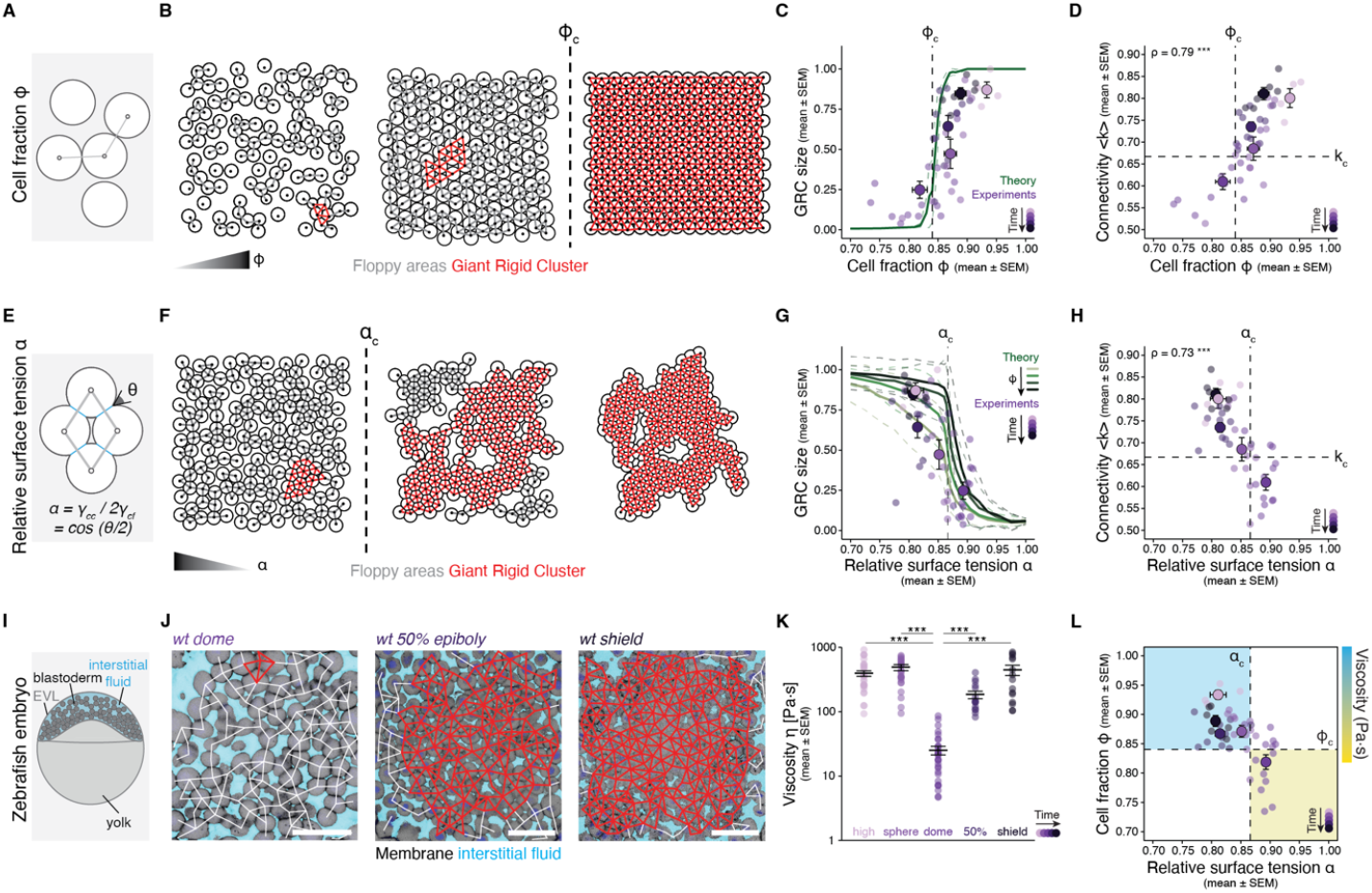
Critical points in cell fraction, contact surface tensions and cell connectivity are crossed simultaneously during a rigidity transition in the zebrafish blastoderm. **(A)** Schematic diagram of density-dependent connectivity. **(B)** Numerical simulations and overlaid connectivity and rigidity maps of jammed hard discs performed using the Lubachevsky-Stillinger algorithm ^34^ at different fraction *ϕ* values. Rigidity emerges at *ϕ*_*c*_ ≈ 0.84. **(C)** Plot of the fraction of the network occupied by the GRC as a function of cell fraction *ϕ* for the simulations shown in (B) with overlaid experimental data from zebrafish embryos from high until shield stage (embryos per stage: high n = 4, sphere n = 11, dome n = 14, 50% epiboly n = 11, shield n = 6). **(D)** Plot of normalized connectivity <*k*> as a function of cell fraction *ϕ* from the experimental data shown in (C). **(E)** Schematic diagram of adhesion-dependent connectivity. Relative surface tension *α* is defined by the Young-Dupré relation and can be measured by the angle θ formed at the contact edge. **(F)** Numerical simulations and overlaid connectivity and rigidity maps of cell arrays with *ϕ* < *ϕ*_*c*_ at different relative surface tension *α* values. Rigidity emerges at *α*_*c*_ ≈ 0.8662. **(G)** Plot of the fraction of the network occupied by the GRC as a function of relative surface tension *α* for the simulations shown in (F), performed for different cell fraction *ϕ* values below *ϕ*_*c*_, and overlaid experimental data from zebrafish embryos from high until shield stage (same as in (C)). **(H)** Plot of normalized connectivity <*k*> as a function of relative surface tension *α* from the experimental data shown in (C). **(I)** Schematic diagram of the zebrafish dome-stage embryo. EVL: enveloping layer. **(J)** Exemplary 2D confocal sections at the 2^nd^ deep-cell layer of the blastoderm overlaid with their connectivity and rigidity maps at dome, 50% epiboly and shield stages. Interstitial fluid is marked with dextran-647, nuclei with H2A-BFP, H2A-mCherry, or H2B-GFP, and membranes with *α*-catenin-citrine or LynTomato. **(K)** Plot of the central blastoderm viscosity values in zebrafish embryos from high to shield stage, as measured via micropipette aspiration ^5^ (embryos per stage: high n = 30, sphere n = 26, dome n = 30, 50% epiboly n = 15, shield n = 16). **(L)** Phase diagram of tissue viscosity with coordinates cell fraction *ϕ* and relative surface tension *α* and overlaid experimental data from (C) and (G) showing that wildtype embryos are constrained to two out of the four regimes. Cyan-shaded areas indicate high viscosity and yellow-shaded areas indicate low viscosity. Floppy areas are illustrated in gray, GRC in red. Light purple color: early stage. Dark purple color: late stage. Dashed lines: critical points. ρ: Spearman correlation (D, H); Kruskal-Wallis with Dunn’s post hoc test (Dunnett adjustment) (K). Scale bars: 50 µm in (J).

The standard theory of jamming transitions predicts that beyond a critical point in cell fraction *ϕ*_*c*_ (*ϕ*_*c*_ ≈ 0.84), the rheological response of the tissue is determined by the fact that cells are *jammed*, meaning that stresses propagate throughout large tissue fractions and any deformation implies an energy penalty ^25,28–30^. When looking at the topology of cell-cell contacts, *ϕ*_*c*_ coincides with the emergence of generic rigidity (**Fig. 1A, B**) ^31^. At *ϕ*_*c*_, the cell-cell contact network crosses the critical point <*k*_*c*_> (<*k*_*c*_>~4 in 2D), leading to rigid cluster percolation (**Fig. 1B, C**) ^22,26^. When quantifying cell fraction, connectivity, GRC ^32,33^ and tissue viscosity in the early zebrafish embryo, we observe that *wildtype* (*wt*) tissues with cell fraction above the critical point display connectivity above the rigidity percolation transition threshold, a large GRC and high viscosity (**Fig. 1C, D, K, Fig. S1B-D**). These results suggest that the material properties of early embryonic tissues follow the physics of density-dependent jamming of granular materials.

Previous reports showed that jamming-dependent rigid cluster percolation appears both when contacts between cells are passive –as in hard, non-attractive or repulsive spheres ^34,35^– or active –as in cell-cell contact networks with active adhesion mechanisms ^19,23,28^. To explore the contribution of active adhesion to rigid cluster percolation we theoretically approached cell-cell adhesive interactions within the soap bubble theoretical framework. We used the non-dimensional parameter *α*, defined as the ratio between cell-cell γ_*cc*_ and cell-fluid γ_*cf*_ surface tensions, which can be estimated from the angle formed at the contact edge between two cells, θ ^19,36–40^ (**Fig. 1E**, see Supplementary Theory Note). To examine how rigid cluster percolation is influenced by the relative surface tension we mathematically explored the behavior of a floppy motif of four cells under different *α* values (**Fig. S1F**). We analytically identified a critical point, *α*_*c*_ ≈ 0.866, at which a new contact is formed and thereby rigidifying the motif (**Fig. S1F**, see Supplementary Theory Note). To address if *α*_*c*_ can explain rigidification in an arbitrary array of cells, we generated *in silico* tilings at the hard-disk regime (*α* = 1, no adhesion) whose cell fraction is below the critical jamming fraction (*ϕ* < *ϕ*_*c*_) and for which we verified that the GRC size is negligible (**Fig. 1F**). Then, we decreased *α* in a quasi-static way using the *Surface Evolver* software ^41^ (see Supplementary Theory Note). Once the system reached the optimal configuration for a given *α*, we computed GRC size. At *α*_*c*_, a jump in the average connectivity above <*k*_*c*_> and a transition in GRC size are observed, with higher initial packing fractions resulting in a sharper transition (**Fig. 1F, G, Fig. S1G**). This shows that rigidification can be driven by only changing contact surface tensions. The parameter *α* therefore acts as a new control parameter for a rigidity transition. Importantly, when quantifying contact surface tensions in zebrafish embryos, we observe that tissues of high viscosity, large GRC, and with connectivity and cell fraction above their corresponding critical points also have *α* below *α*_*c*_ (**Fig. 1H, Fig. S1E**). This suggests that early embryonic tissues can also undergo adhesion-dependent jamming.

Based on these findings, we propose a 2D (*α, ϕ*) phase diagram of the tissue material state organized around the critical points in relative surface tension *α*_*c*_ and cell fraction *ϕ*_*c*_ (**Fig. 1L**). *wt* embryos are mostly constrained to two of the four potential material regimes: the upper left regime, corresponding to topologically rigid tissues of high viscosity, and the bottom right regime, corresponding to topologically floppy tissues of low viscosity (**Fig. 1L**). During this transition early embryos cross simultaneously the critical points *ϕ*_*c*_, *α*_*c*_ and <*k*_*c*_> by coupling changes in cell fraction, contact surface tensions and cell connectivity. This raises the question if these control parameters can be uncoupled and how each parameter contributes to the tissue rheological state.

### Relative surface tension *α* is the major control parameter of the tissue material state *in vivo*

To address the above questions, we experimentally uncoupled cell fraction, relative surface tension and connectivity and measured tissue viscosity using micropipette aspiration ^5,24^. We followed two approaches: we used (i) a mutant line *MZpky* ^42^, in which the outermost epithelium is disrupted and the interstitial fluid leaks out, leading to tissues of higher cell fraction (**Fig. 2A, B, Fig. S2A**); and (ii) zebrafish embryonic explants ^43,44^, which at shield stage accumulate more interstitial fluid, displaying lower cell fraction (**Fig. 2C, D, Fig. S2A**). Intriguingly, both manipulations escape the constraints that couple fraction and contact surface tensions in *wt* embryos. *MZpky* embryos have cell fraction above *ϕ*_*c*_, connectivity above <*k*_*c*_> and relative surface tension above *α*_*c*_, falling in the upper right regime of the phase diagram, whereas zebrafish explants have cell fraction below *ϕ*_*c*_, connectivity above <*k*_*c*_ > and relative surface tension below *α*_*c*_, falling in the lower left regime of the phase diagram (**Fig. 2E, Fig. S2A, B, E**). Surprisingly, *MZpky* embryos display low viscosity despite appearing topologically rigid, and, conversely, zebrafish explants are topologically rigid and highly viscous despite having cell fraction below *ϕ*_*c*_ (**Fig. 2F, G, Fig. S2C, D**). These results show that when cell density and adhesion changes are uncoupled, the tissue material state is determined by cell-cell adhesion strength.

**Figure 2:**
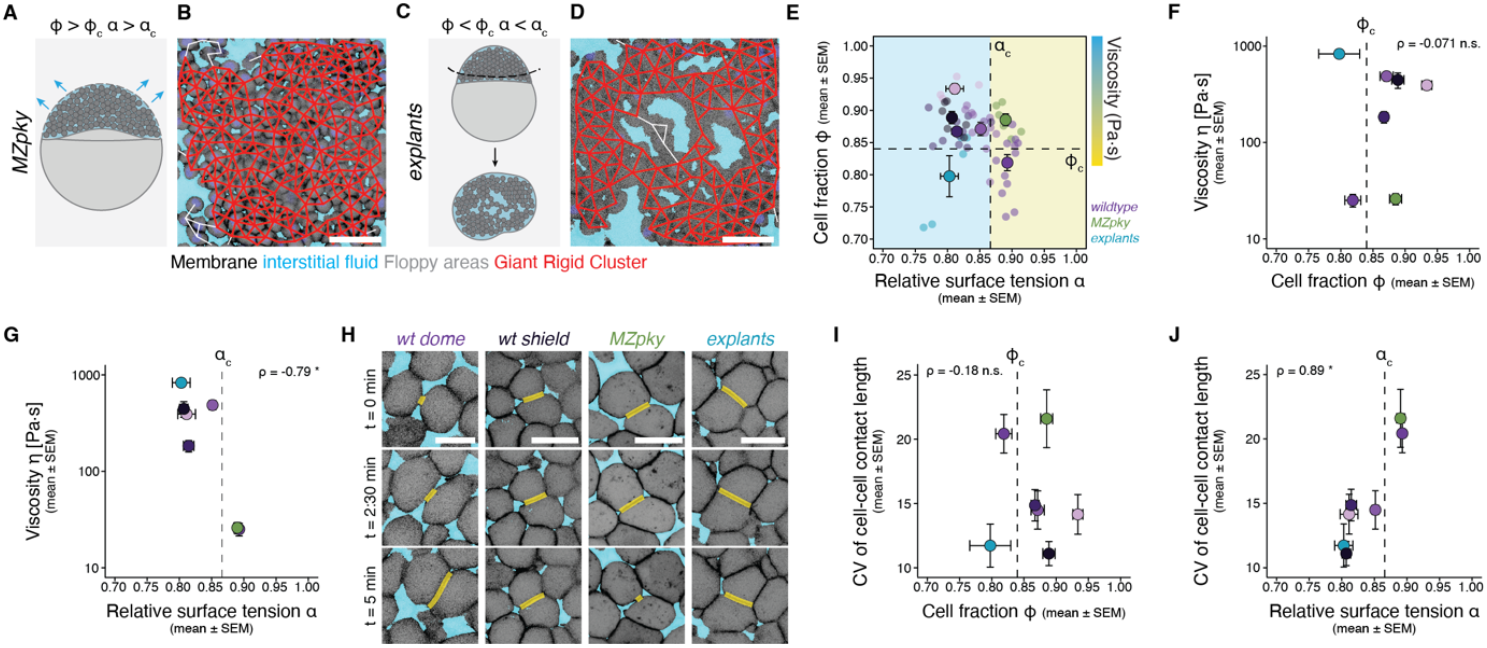
Decoupling the contributions of cell fraction and contact surface tensions to the tissue material state. **(A)** Schematic diagram and **(B)** exemplary 2D confocal section at the 2^nd^ deep-cell layer of the blastoderm of *MZpky* embryos, in which the interstitial fluid leaks out. Membranes labeled with membrane-GFP, nuclei with H2A-BFP, and interstitial fluid with dextran-647. **(C)** Schematic diagram and **(D)** exemplary 2D confocal section of the deep-cell tissue of blastoderm explants. Membranes labeled with *α*-catenin-citrine, nuclei with H2A-mCherry, and interstitial fluid with dextran-647. **(E)** Phase diagram of tissue viscosity with coordinates cell fraction *ϕ* and relative surface tension *α* and overlaid experimental data from *wt, MZpky* and explanted blastoderms (n = 6 *MZpky* embryos, n = 5 explants; *wt* as in Fig. 1L). Cyan-shaded areas indicate high viscosity and yellow-shaded areas indicate low viscosity. **(F-G)** Plots of blastoderm viscosity values for the conditions shown in (E), as a function of cell fraction *ϕ* (F) and relative surface tension *α* (G) (n = 16 *MZpky* and n = 6 explants). **(H)** Representative high-resolution 2D confocal sections from time-lapse imaging over 10 min at 10-sec intervals (images shown at t = 0, 150, and 300 sec) of *wt* dome and shield stage, *MZpky*, and explants. The yellow-shaded line marks the cell-cell contact length. Membranes labeled with *α*-catenin-citrine or membrane-GFP, interstitial fluid with dextran-647. **(I-J)** Plots of the coefficient of variation (CV) in cell-cell contact length for the conditions shown in (H) as a function of cell fraction *ϕ* (I) and relative surface tension *α* (J) (embryos N and contacts n per stage and condition: *wt* high N = 3, n = 13; *wt* sphere N = 3, n = 12; wt dome N = 8, n = 39; *wt* 50% epiboly N = 7, n = 19; *wt* germ ring to shield N = 8, n = 36; *MZpky* N = 6, n = 27; explants N = 5, n = 21). Dashed lines: critical points. ρ: Spearman correlation (F, G, I, J). Scale bars: 50 µm in (B) and (D), 20 µm in (H).

To understand why relative surface tension is a better predictor of the tissue material state than cell fraction, we examined its relationship with active cell parameters, such as the previously reported cell-cell contact length fluctuations (CLFs) ^18^ (**Fig. 2H, Fig. S2F**). We observe a dependency between relative surface tension, CLFs and tissue viscosity, where all tissues above *α*_*c*_ display large CLFs and are fluidized (**Fig. 2J**), irrespective of their cell fraction (**Fig. 2I**). This suggests that even when *ϕ* > *ϕ*_*c*_ and <*k*> > *k*_*c*_, cell-cell contacts are highly dynamic when *α* > *α*_*c*_, allowing the tissue to deform, explaining why relative surface tension *α* is an accurate proxy for the tissue material response. These results show that although early embryonic tissues couple density-dependent and adhesion/CLF-dependent material phase transitions, their material state is only determined by the latter, raising the question of the function of control parameter coupling in morphogenesis.

### Precise engineering of tissue material states via uncoupling the multiple control parameters

To explore if and how coupling of the control parameters impacts the tissue morphogenetic outcome, we engineered the macroscopic material state (fluid-like vs. solid-like) of *wt* embryos by either coupling or uncoupling the changes in cell fraction and relative surface tension. To do this, we combined several methodologies to precisely fine-tune them in relation to their corresponding critical points. We performed experimental manipulations in: i) *wt* dome-stage embryos, which have high relative surface tension (*α* > *α*_*c*_), low cell fraction (*ϕ* < *ϕ*_*c*_) and are topologically floppy (small GRC, <*k*> < *k*_*c*_) and rheologically fluid-like (large CLFs), and ii) *wt* 50% epiboly stage embryos, which have low relative surface tension (*α* < *α*_*c*_), high cell fraction (*ϕ* > *ϕ*_*c*_) and are topologically rigid (big GRC, < *k* > > *k*_*c*_) and rheologically solid-like (small CLFs) (**Fig. S3A-E**, light orange vs. light pink data points).

To change cell fraction, we interfered with osmosis ^45^. To increase cell fraction, we induced a hypotonic environment by using ouabain, a Na^+^/K^+^-ATPase inhibitor, whereas to decrease cell fraction we induced a hypertonic environment by injecting the sugar alcohol mannitol in the interstitial fluid ^46^. This approach in dome-stage embryos fine-tunes cell fraction above or below the critical *ϕ*_*c*_ (**Fig. S3A**, bright green vs. dark orange data points), resulting in populations occupying the upper right and lower right regimes of the phase diagram, respectively (**Fig. 3A**).

**Figure 3:**
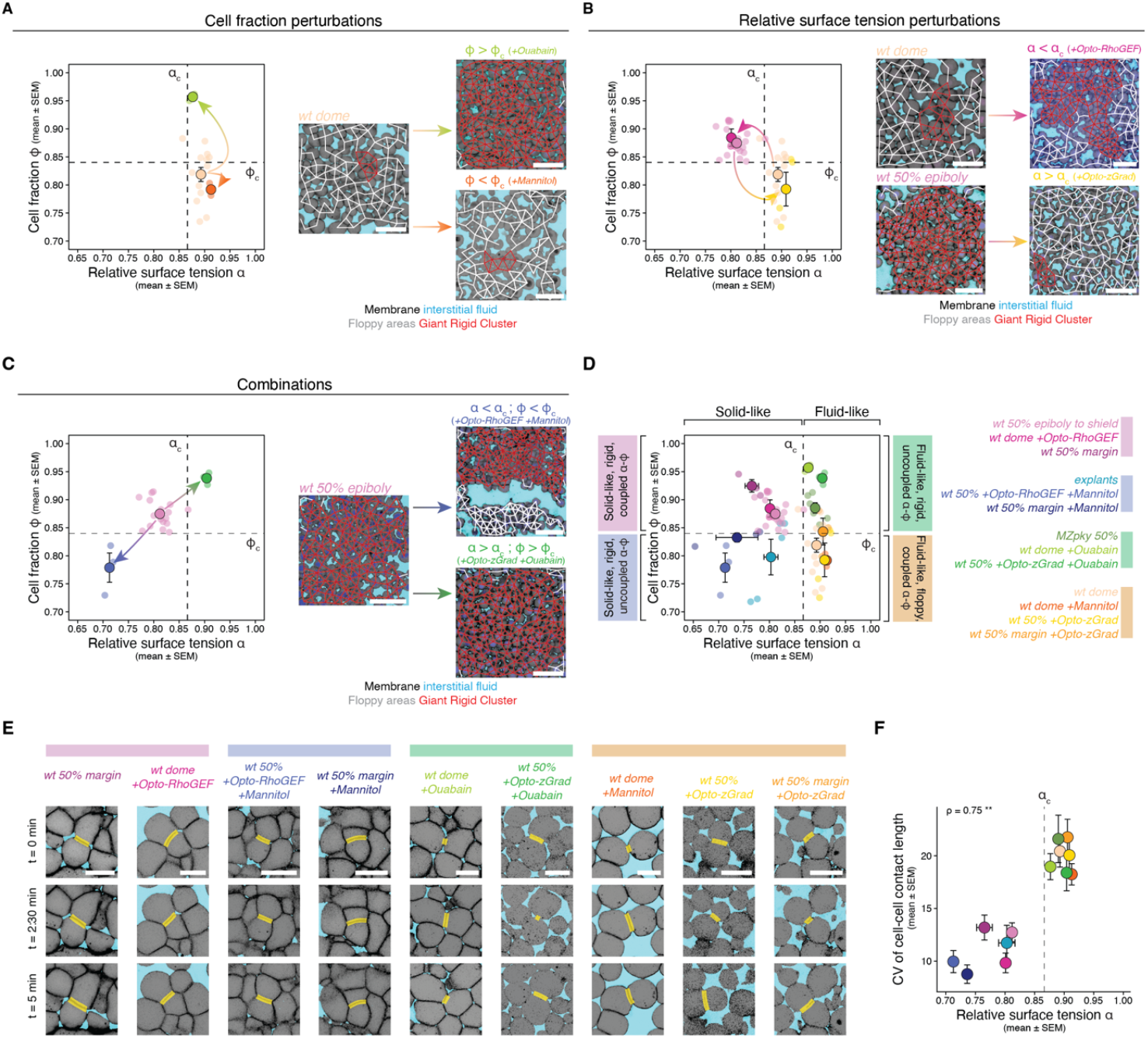
Precise engineering of tissue material states via independent fine-tuning of the control parameters. **(A)** Phase diagram of the cell fraction perturbation experiments, with coordinates cell fraction *ϕ* and relative surface tension *α* (left) and representative 2D confocal sections of the blastoderm with overlaid connectivity and rigidity maps (right) from dome-stage embryos, either untreated (*wt*) or osmotically altered with ouabain or mannitol to increase or decrease cell fraction in relation to *ϕ*_*c*_ (embryo numbers as in (D)). **(B)** Phase diagram with the parameters shown in (A) (left) and representative 2D confocal sections of the blastoderm with overlaid connectivity and rigidity maps (right) from *wt* or optogenetically rigidified (Opto-RhoGEF) dome-stage embryos and from *wt* or optogenetically fluidized (Opto-zGrad) 50% epiboly-stage embryos (embryo numbers as in (D)). **(C)** Phase diagram with the parameters shown in (A) (left) and representative 2D confocal sections of the blastoderm with overlaid connectivity and rigidity maps (right) from 50% epiboly-stage embryos, either *wt* or optogenetically rigidified in a hypertonic solution (Opto-RhoGEF + Mannitol) or optogenetically fluidized in a hypotonic solution (Opto-zGrad + Ouabain) (embryo numbers as in (D)). **(D)** Phase diagram with the parameters shown in (A) with overlaid experimental data from different combinations of genetic, optogenetic, and osmotic manipulations occupying the four material regimes of the phase diagram (embryo numbers: *wt* 50% epiboly to shield n = 17, explants n = 5, *MZpky* n = 6, *wt* dome n = 14, *wt* 50% + opto-zGrad n = 4, and n = 3 embryos for *wt* dome + opto-RhoGEF, *wt* 50% margin, *wt* 50% + opto-RhoGEF + mannitol, *wt* 50% margin + mannitol, *wt* dome + ouabain, *wt* 50% + opto-zGrad + ouabain, *wt* dome + mannitol, and *wt* 50% margin + opto-zGrad). **(E)** Representative high-resolution 2D confocal sections from time-lapse imaging over 10 min at 10-sec intervals (images shown at t = 0, 150, and 300 sec) of the experimental conditions shown in (D). The yellow-shaded line marks the cell-cell contact length. **(F)** Plots of the coefficient of variation (CV) in cell-cell contact length for the conditions shown in (E) as a function of relative surface tension *α* (embryo numbers for *α* as in (D); embryos N and contacts n for contact length fluctuations per stage and condition: *wt* 50% epiboly to shield N = 13, n = 45; *wt* dome + opto-RhoGEF N = 4, n = 18; *wt* 50% margin N = 4, n = 15; explants N = 5, n = 21; *wt* 50% + opto-RhoGEF + mannitol N = 3, n = 15; *wt* 50% margin + mannitol N = 3, n = 20; *MZpky* N = 6, n = 27; *wt* dome + ouabain N = 5, n = 35; *wt* 50% + opto-zGrad + ouabain N = 3, n = 15; *wt* dome N = 8, n = 39; *wt* dome + mannitol N = 6, n = 30; *wt* 50% + opto-zGrad N = 5, n = 20; *wt* 50% margin + opto-zGrad N = 3, n = 15). Dashed lines: critical points. ρ: Spearman correlation (F). Membranes labeled with *α*-catenin-citrine or LynTomato, nuclei with H2A-BFP, H2B-GFP, or H2A-mCherry, and interstitial fluid with dextran-647. Scale bars: 50 µm in (A), (B) and (C), 20 µm in (E).

To change cell-cell adhesion strength, we established two optogenetic tools acting on the relative surface tension. In agreement with previous *ex vivo* assays ^36,47^, we find that the relative surface tension *α* in the blastoderm cells *in vivo* arises from the differential localization of adherens junction components at the cell-cell contact interface and of active myosin at the cell-fluid interface (**Fig. S3G**, red vs. cyan arrowheads). To decrease *α*, we increased γ_*cf*_ by enhancing myosin activity (**Fig. S3H**), whereas to increase *α* we increased γ_*cc*_ by reducing *α*-catenin protein levels (**Fig. S3I**), using optogenetics to precisely fine-tune *α* just above or just below the critical point. For instance, to decrease *α* and rigidify *wt* dome-stage embryos, we used the opto-RhoGEF tool, which sequesters RhoGEF to the plasma membrane upon light activation ^48^ (**Fig. 3B, Fig. S3H-H’’’, J, L**, see Methods). In contrast, to increase *α* and fluidize *wt* 50% epiboly-stage embryos, we generated the opto-zGrad tool, which degrades *α*-catenin upon light exposure (**Fig. 3B, Fig. S3I, K-M**, see Methods). The opto-zGrad tool is a nanobody-based degron system targeting GFP and its variants, combined with an optogenetic module, AsLOV2, which enables light-controlled protein binding ^49,50^ (**Fig. S3I)**. We expressed opto-zGrad in a zebrafish knock-in line for *α*-catenin fused to citrine ^51^ to induce *α*-catenin degradation, weaken cell-cell adhesion and thus increase the relative surface tension *α* and induce tissue fluidization (**Fig. S3A-D**). Given that in these manipulations cell fraction also changes (**Fig. 3B, Fig. S3A, B**), we uncoupled the two parameters using combinations of the above tools. For instance, optogenetic decrease of *α* using opto-RhoGEF also increased cell fraction (**Fig. 3B, Fig. S3A**), but combination with mannitol treatment led to the generation of tissues occupying the bottom left regime of the phase diagram (**Fig. 3C**, pink-to-blue arrow, **Fig. S3A, B**). Additionally, optogenetic increase of *α* using opto-zGrad also reduced cell fraction (**Fig. 3B, Fig. S3A**), however when combined with ouabain treatment resulted in tissues occupying the upper right regime of the phase diagram (**Fig. 3C**, pink-to-green arrow, **Fig. S3A, B**). By expanding these manipulations to the marginal region of the blastoderm, which is topologically rigid and highly viscous ^5,19^, we managed to start from the pluripotent zebrafish blastoderm and engineered it to occupy all the predicted regimes of the phase diagram via a variety of experimental approaches that fine-tune the control parameters with high precision (**Fig. 3D, Fig. S3A, B**).

Having this collection of tissues, we then asked if cell fraction and relative surface tension are sufficient parameters to predict material states. To this end, we quantified connectivity (**Fig. S3C**) and rigid cluster percolation (**Fig. S3D**) to map topological rigidity, and CLFs to estimate tissue viscosity (**Fig. 3E, Fig. S3E**). We find that all engineered tissues with *α* < *α*_*c*_ are solid-like (low CLFs) and topologically rigid (<*k*> > *k*_*c*_, large GRC) irrespective of their cell fraction (**Fig. 3F, Fig. S3F**). Conversely, although all engineered tissues with *α* > *α*_*c*_ are fluid-like (high CLFs), they are topologically floppy only when cell fraction is *ϕ* < *ϕ*_*c*_. This leads to two material states, a solid-like and a fluid-like, where each one can be further separated into two regimes depending on whether they couple changes in *α* and *ϕ* (**Fig. 3D**): (i) upper left: solid-like, rigid, coupled *α*-*ϕ*; (ii) bottom left: solid-like, rigid, uncoupled *α*-*ϕ*; (iii) upper right: fluid-like, rigid, uncoupled *α*-*ϕ*; and (iv) bottom right: fluid-like, floppy, coupled *α*-*ϕ*. Thus, tuning solely relative surface tension and cell fraction in relation to their critical points, *α*_*c*_ and *ϕ*_*c*_, is sufficient to engineer tissues with accurately controlled material states.

### The tissue material state discretizes tissue organization via formation of tricellular contacts

The above engineered tissues allow us to explore how the tissue material state and the coupling of the underlying control parameters impacts the morphogenetic outcome. To quantify morphogenetic outcomes, we first performed a quantitative morphological analysis of the engineered tissues shown in Fig. 3D using a custom-made image analysis pipeline for extraction of morphological measurements (see Methods) (**Fig. 4A**). We then performed dimensionality reduction using principal component analysis (PCA) on eight morpho-features associated with cell shape, cell surface and the extracellular space (**Fig. 4A**, see Methods), without including control and order parameters of the tissue material state (*α, ϕ*, <*k*>, GRC, CLFs). This allowed us to represent the tissues as points in a low-dimensional morphospace, with axes determined by the PCs (**Fig. 4B)**. We find that PC1 and PC2 accounted for 63.9% and 16.2% of the variance in the data, respectively, with little overlap between fluid-like and solid-like tissues. Importantly, analyzing the relationship between the PCs and the control parameters revealed that PC1 changes as a function of the relative surface tension *α* whereas PC2 does so as a function of the cell fraction *ϕ* (**Fig. 4C, Fig. S4B**), suggesting that the tissue material state but also the control parameters used are responsible for generating the major morphological differences in tissue organization.

**Figure 4:**
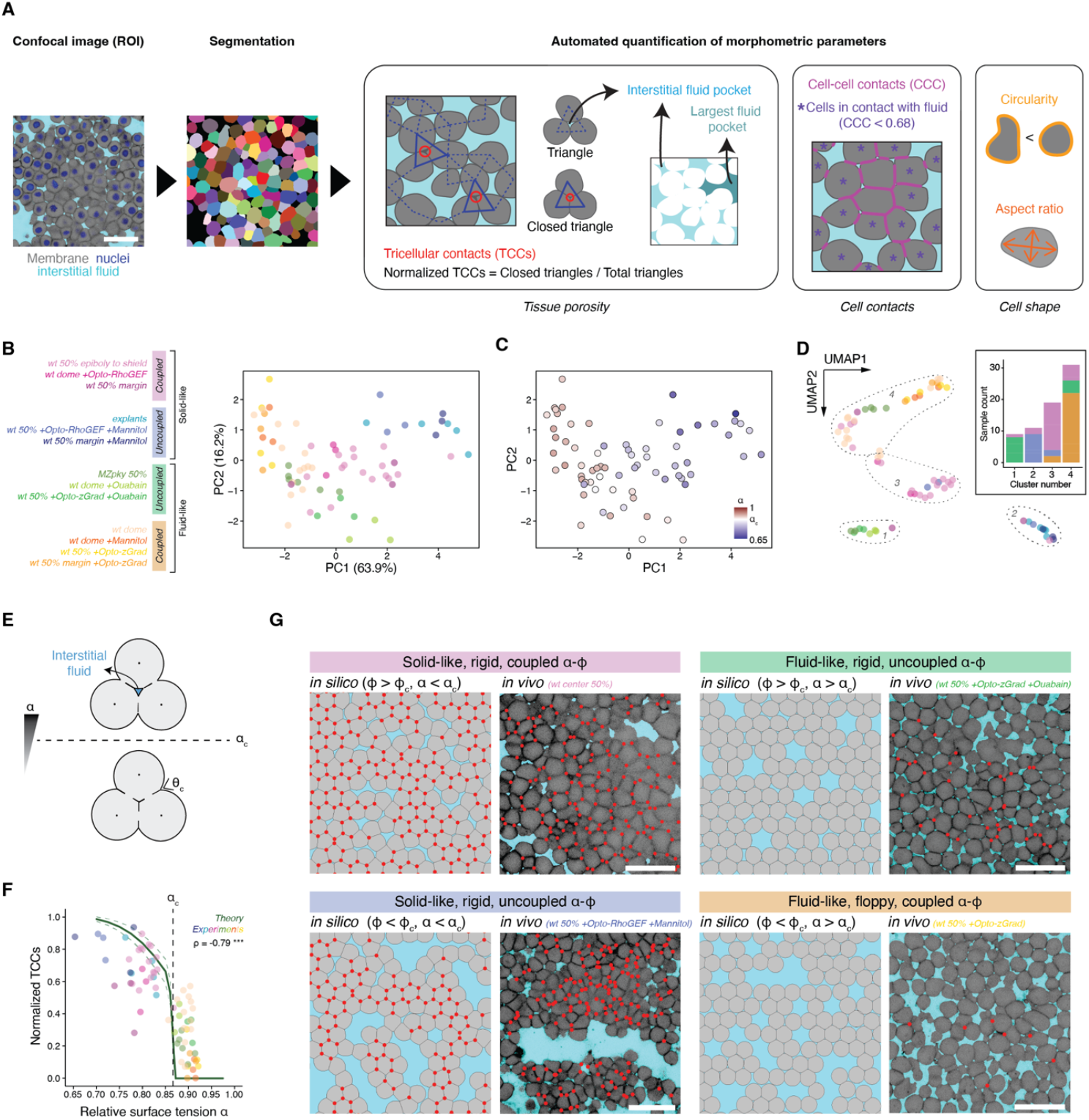
Contact surface tension-driven rigidification leads to abrupt formation of tricellular contacts. **(A)** Schematic diagram of the image analysis pipeline used for cell segmentation and morphometric analysis. **(B)** PCA plot of the morphology of tissues occupying the material phase diagram in Fig. 3D. **(C)** Same PCA plot as in (B), color-coded according to the relative surface tension *α* values for each tissue. **(D)** UMAP plot of the morphology of tissues occupying the material phase diagram in Fig. 3D, with the clusters identified with k-means clustering outlined with dashed lines, and a histogram of the tissue material phase regime composition of each cluster. **(E)** Representation of the computation of *α*_*c*_ at which interstitial spaces between three cells are closed (see Supplementary Theory Note). **(F)** Simulation at the tissue-level scale computing the fraction of TCCs that are closed as a function of *α*, with overlaid experimental values from all the tissues shown in (B). Dashed line: critical point. **(G)** Exemplary tissue-level simulations (left) and 2D confocal sections of experimental tissues for the four material regimes. The red dots indicate closed TCCs. Membranes labeled with *α*-catenin-citrine, interstitial fluid with dextran-647. ρ: Spearman correlation for experimental data (F). Scale bars: 50 µm.

Intriguingly, although the engineered tissues belonging to the same material regime exhibit a continuum of values in relative surface tension *α* and cell fraction *ϕ*, at the morphological level they exhibit discrete tissue architectures according to their material regime (**Fig. S4A**). We applied Uniform Manifold Approximation and Projection (UMAP) to our dataset as a non-linear dimensionality reduction technique ^52^, followed by k-means clustering to group the datapoints based on similarities in their morpho-features ^53,54^. Intriguingly, the four morphological clusters appear to correlate with the four material regimes (**Fig. 4D**): Clusters 1 and 4 consist almost exclusively of data points corresponding to fluid-like tissues (*α* > *α*_*c*_) and clusters 2 and 3 of data points corresponding to solid-like tissues (*α* < *α*_*c*_). Additionally, there is a clear separation between the coupled vs. uncoupled regimes (for fluid-like tissues: cluster 4, coupled *α*-*ϕ*; cluster 1, uncoupled *α*-*ϕ*; for solid-like tissues: cluster 3, coupled *α*-*ϕ*; cluster 2, uncoupled *α*-*ϕ*). This suggests that the critical points defining the material regimes are also defining the limits of their morphological regimes, discretizing the potential tissue organization states a tissue can exhibit.

To identify the key dimension that accounts for the major morphological differences, we examined which morpho-features primarily contribute to the PCs and found that PC1 is strongly correlated with features related to the formation of tricellular contacts (TCCs). In triangular cell motifs, we find that the central interstitial space vanishes and the cells form a TCC (**Fig. 4E**) depending on the relative surface tension, in agreement with previous work ^18,55^. Intriguingly, the formation of the TCC occurs exactly at the critical point *α*_*c*_ ≈ 0.866, for which we have shown that tissue rigidification takes place (see Supplementary Theory Note). At the collective level, a sharp increase in TCCs at *α*_*c*_ is observed in arbitrarily disordered cell packings (**Fig. 4F, G**), unfolding the potentiality of being fully confluent and thus defining a transition from porous to non-porous regime (see Supplementary Theory Note). Importantly, quantification of the TCCs in our experimental data revealed that solid-like tissues (*α* < *α*_*c*_) display a high number of TCCs, whereas fluid-like tissues (*α* > *α*_*c*_) display very few TCCs, independently of their cell fraction (**Fig. 4F, G, Fig. S4C**). Similarly, tissue porosity changes drastically at *α*_*c*_, but not at *ϕ*_*c*_ (**Fig. S4D, E**), showing that adhesion-driven rigidity transitions are coupled to porosity transitions. Altogether, these results show that the tissue material state dictates tissue morphology and organization via the formation of TCCs and a concomitant loss of porosity.

### Adhesion-driven solidification triggers apicobasal polarity in pluripotent tissues

Tissue architecture has been recently found to be an important modulator of cell decisionmaking ^56^. Our results, which pinpoint the tissue material state as a significant factor of tissue organization, raise the question of whether amorphous and still pluripotent tissues can steer their fate potential based on the type of phase transition they undergo. Specifically, the abrupt formation of TCCs resembles the tricellular junctions (TCJs) observed in epithelia, which are high-tension specialized structures characterized by the presence of specific proteins such as Tricellulin/MARVELD2 and Angulin-1/LSR ^57–60^. To address if the TCCs observed above resemble true TCJs, we performed a quantitative analysis of their protein composition via live imaging. We found that pluripotent tissues undergoing adhesion-driven solidification accumulate the TCJ-specific proteins Marveld2a and Lsr and the mechanosensitive proteins F-actin and *α*-catenin, but are depleted of the adherens junction protein E-cadherin (**Fig. S5A-E**), suggesting that adhesion-driven rigidification in undifferentiated tissues may prompt them to acquire epithelial character. This further raises the question of how the coupling with the other control parameters may contribute to the epithelial fate potential.

To explore this, we performed longer time lapses of the coupled vs. uncoupled solidification (**Fig. 5A**). This revealed that, although in both cases rigidification is accompanied by closure of the TCJs and local exclusion of interstitial fluid from between the cells, in the coupled solid-like regime the interstitial fluid is redistributed in small pockets at regions that have not yet closed the tricellular gaps. In contrast, in all experimental conditions in the uncoupled solid-like regime, the interstitial fluid is redistributed in larger pockets that eventually coalesce into bigger lumen-like structures (**Fig. 5B, Fig. S4A**). Given that lumen formation is often accompanied by the acquisition of apicobasal polarity ^61^, we performed immunostaining of several polarity markers. Strikingly, we observed that uncoupled solidification not only leads to morphological luminogenesis, but also to the acquisition of apicobasal polarity, as observed by the absence of E-cadherin and enrichment of F-actin, P-MLC, and ZO-1 at the interface with the lumen (**Fig. 5D, E**). This shows that adhesion-driven rigidity favors epithelial-like character via the formation of TCJs and loss of porosity, and when the transition occurs via uncoupling changes in contact surface tension and fraction, it further promotes epithelial organization via lumen formation and acquisition of apicobasal polarity at the molecular level.

**Figure 5:**
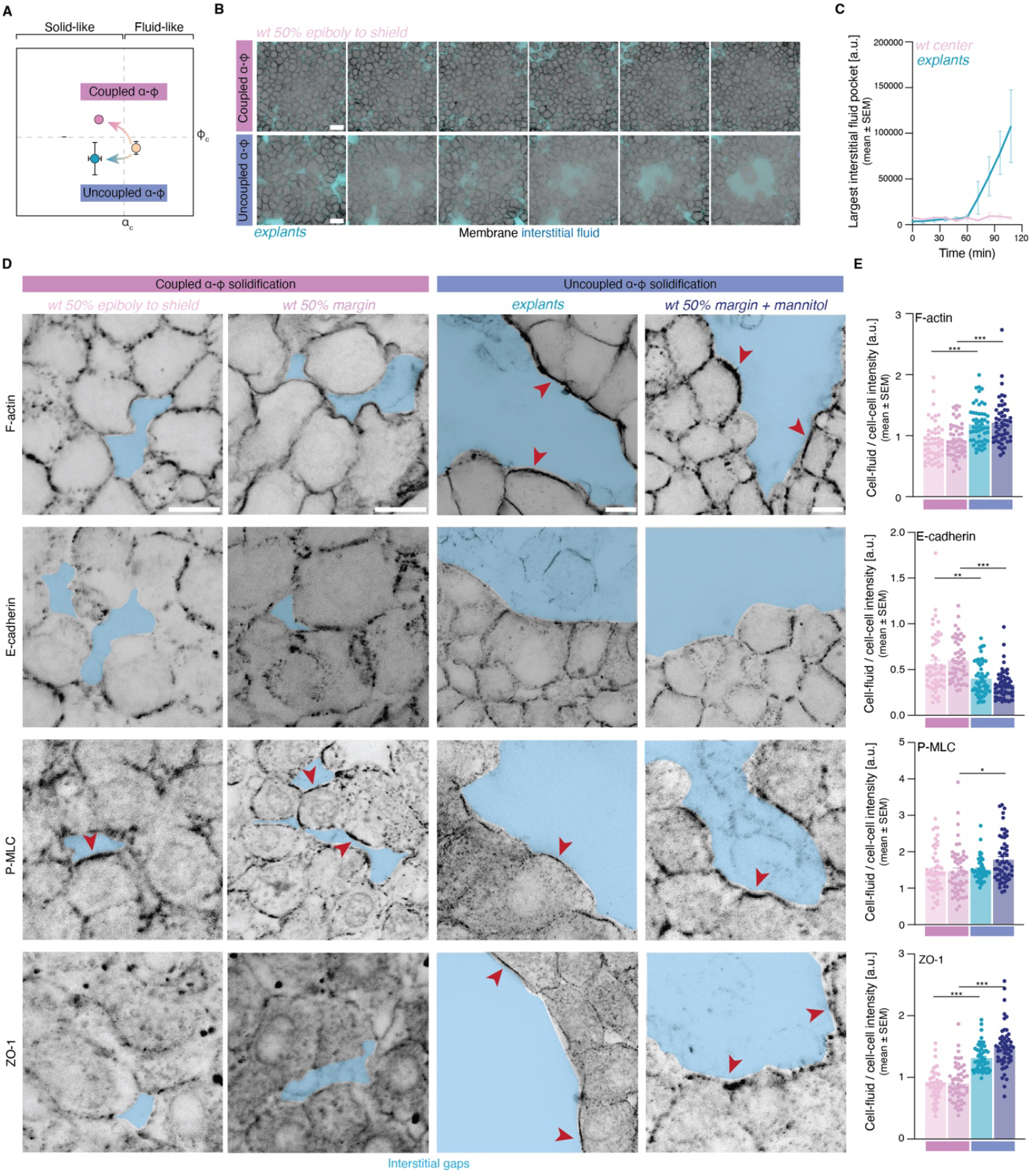
Uncoupling density-from adhesion-dependent solidification in pluripotent tissues triggers the acquisition of epithelial polarity. **(A)** Phase diagram of the tissue material regimes as a function of cell fraction *ϕ* and relative surface tension *α* showcasing tissue solidification by coupling (magenta) vs. uncoupling (blue) the two control parameters. **(B)** Time stills (12 min interval) from live imaging of coupled (*wt* center) vs. uncoupled (explants) tissue solidification, showing lumen formation in the latter. **(C)** Plot of the quantification of the number of pixels in the largest interstitial fluid pocket over time for the experimental conditions in (B) (*wt* N = 2, explants N = 3). **(D)** Exemplary high resolution 2D confocal sections of samples undergoing coupled (*wt* center and margin) vs. uncoupled (explants and *wt* margin + mannitol) tissue solidification and stained for F-actin (phalloidin), E-cadherin, P-MLC and ZO-1. **(E)** Plots of quantifications of the intensity for the proteins shown in (D) at the cell-interstitial fluid vs. cell-cell interfaces (n = 50 cell-cell and cell-fluid surfaces, N = 4 embryos for each staining and each experimental condition). Interstitial fluid manually labeled in light blue. Red arrowheads indicate sites of protein accumulation. Kruskal-Wallis test (E). Scale bars: 30 µm (B), 10 µm (D).

### OUTLOOK

This work shows that the dynamic co-regulation and fine-tuning of the multiple microscopic control parameters in living materials not only sets their macroscopic state as a binary switch, as expected from inert glassy systems, but also their morphogenetic potential. We focused on the roles of density and adhesion, proposing a new phase diagram for living tissues deviating from the classic diagrams proposed for inert materials ^29^. In non-confluent pluripotent tissues, solidification is driven by an adhesion-driven rigidity transition, which is accompanied by a sudden formation of tricellular junctions, loss of porosity and reduction of contact dynamics. Intriguingly, if adhesion-driven rigidification occurs independently of density-driven rigidification, tissues form lumina and eventually acquire apicobasal polarity. Tricellular junctions, lumina and apicobasal polarity are hallmarks of epithelia, which control cell signaling, e.g. the concentration of diffusive molecules or nutrients ^62–64^, and processes like cell division orientation ^65,66^. Our findings pose tissue phase transitions as part of the instructive cues regulating epithelial organization, and depending on the coupling of the control parameters, tissues acquire different degrees of epithelial character. As a result, a fine co-regulation of the coupling of the control parameters may underlie the diversity of organization states an epithelial tissue can exhibit during development, from simple transient structures, such as the neural tube ^67^ or the Kupffer’s vesicle ^68^, to more complex stable structures, like kidney tubules and mammary gland branches ^69^. Future work linking recent findings on how cell and nucleus shape impact transcriptional dynamics and chromatin modifications ^70–72^ to the coupling of phase transition control parameters could shed light on the molecular mechanisms of rigidification-dependent epithelial organization. We therefore conclude that phase transitions play novel, unexpected roles in developing systems, extending beyond tissue deformability, originating from the ability of living materials to dynamically fine-tune multiple critical points ^73^, which may be a novel mechanism channeling the possible morphogenetic trajectories of developing tissues.

## ACKNOWLEDGEMENTS

We thank Anna Erzberger, Takashi Hiiragi, Daniel Rodríguez Amor, Hanna Engelke, Jared E. Toettcher and members of the Petridou and Corominas-Murtra groups for technical advice, critical discussions and feedback on the manuscript; and the Advanced Light Microscopy Facility (ALMF) and the zebrafish facility at the European Molecular Biology Laboratory (EMBL) for continuous support. We thank Stefano de Renzis for kindly providing initial plasmids for generating the optogenetic constructs and the Heisenberg lab for providing fish lines and plasmids. This work was supported by the Weave project “Tissue material phase transitions and their role in embryo pattern formation” from the Deutsche Forschungsgemeinschaft (DFG, German Research Foundation, 518354236, PE 3800/1-1) to N.I.P. and the Österreichischer Wissenschaftsfonds (FWF, Austrian Science Fund, I6533) to B.C-M. B.C-M. and A.A-T. acknowledge the support of the field of excellence “Complexity of life in basic research and innovation” of the University of Graz.

## AUTHOR CONTRIBUTIONS

N.I.P. and B.C-M. designed the research. L.R-C., C.P-C, and N.I.P. performed the experiments and analyzed the experimental data. L.R-C., C.P-C., and A.A-T. analyzed the tricellular junction data. M.H. and C.A. developed the image morpho-feature analysis pipeline. C.A. and A.M.K. developed the pipeline for reconstruction of connectivity networks. B.C-M. developed all the theoretical models. A.A-T., E.F., and B.C-M. performed the simulations. N.I.P. and B.C-M. wrote the manuscript.

## COMPETING INTERESTS

The authors declare no competing interests.

## METHODS

### Zebrafish handling

Zebrafish (*Danio rerio*) were raised at 28.5°C under a 14 h light / 10 h dark cycle ^74^. The following zebrafish strains were used in this study: wildtype (*wt*) AB, *MZpky* ^75^, *Gt(ctnna-citrine)*_*ct3a 51*_, *Tg(actb2:Lyn-Tomato)* ^76^ and *Tg(actb2:Lifeact-eGFP)* ^77^. Zebrafish embryos were grown at 28.5°C and maintained in 1x Danieau’s medium for all experimental incubations. Staging was performed as previously described ^78^. All animal experiments were carried out according to the guidelines of the Committee for Animal Welfare and Institutional Animal Care and Use (IACUC) under EMBL’s Policy on the Protection and Welfare of Animals Used for Scientific purposes (IACUC code 21-010_HD_NP).

### Cloning

The pCS2(+) vector was used for cloning and mRNA production in all experiments. To generate the opto-zGrad plasmid, two previously described systems were combined: zGrad ^49^ and Opto-Nanobody ^50^. PCR-amplified DNA containing the *short AsLOV2* domain (Addgene Plasmid #159592) and *zGrad* (Addgene Plasmid #119716) was subcloned into a linearized pCS2(+) plasmid (StuI digested and CIP treated) using Gibson Assembly. *AsLOV2* was inserted into the *VHHGFP4* nanobody (used for zGrad; <u>EBI entry</u>) at the outer loop position equivalent to AK74 of LaM8 and LaG9, as described in ^50^. For the opto-RhoGEF tool, opto-RhoAGEF2-mCherry and opto-RhoAGEF2 were subcloned from the ARHGEF11(DHPH)-CRY2-mCherry construct (Addgene Plasmid #89481) into a linearized pCS2(+) vector, using the standard Gibson Assembly protocol. All CIBN constructs, including CIBN-mCherry-CAAX and CIBN-CAAX, were subcloned or engineered from existing constructs in the De Renzis laboratory ^79^. To visualize tricellular junctions, Lsr and Marveld2*a* were selected based on their expression in early developmental stages ^80,81^. The cDNA of *lsr-201* and *marveld2a-202* was cloned from 24 hpf larvae following standard protocols, including RNA extraction using TRIzol™ LS and first-strand cDNA synthesis with the SuperScript® III First-Strand Synthesis System (Invitrogen, Cat. No. 18080051). The cloned fragments were inserted into a pCR™ 2.1-TOPO™ TA vector using the TOPO™ TA Cloning™ Kit (Invitrogen, Cat. No. 450641). Lsr-GFP and Marveld2a-GFP constructs were subsequently generated via Gibson Assembly. The sequences of all oligos used for cloning are provided in the below table.

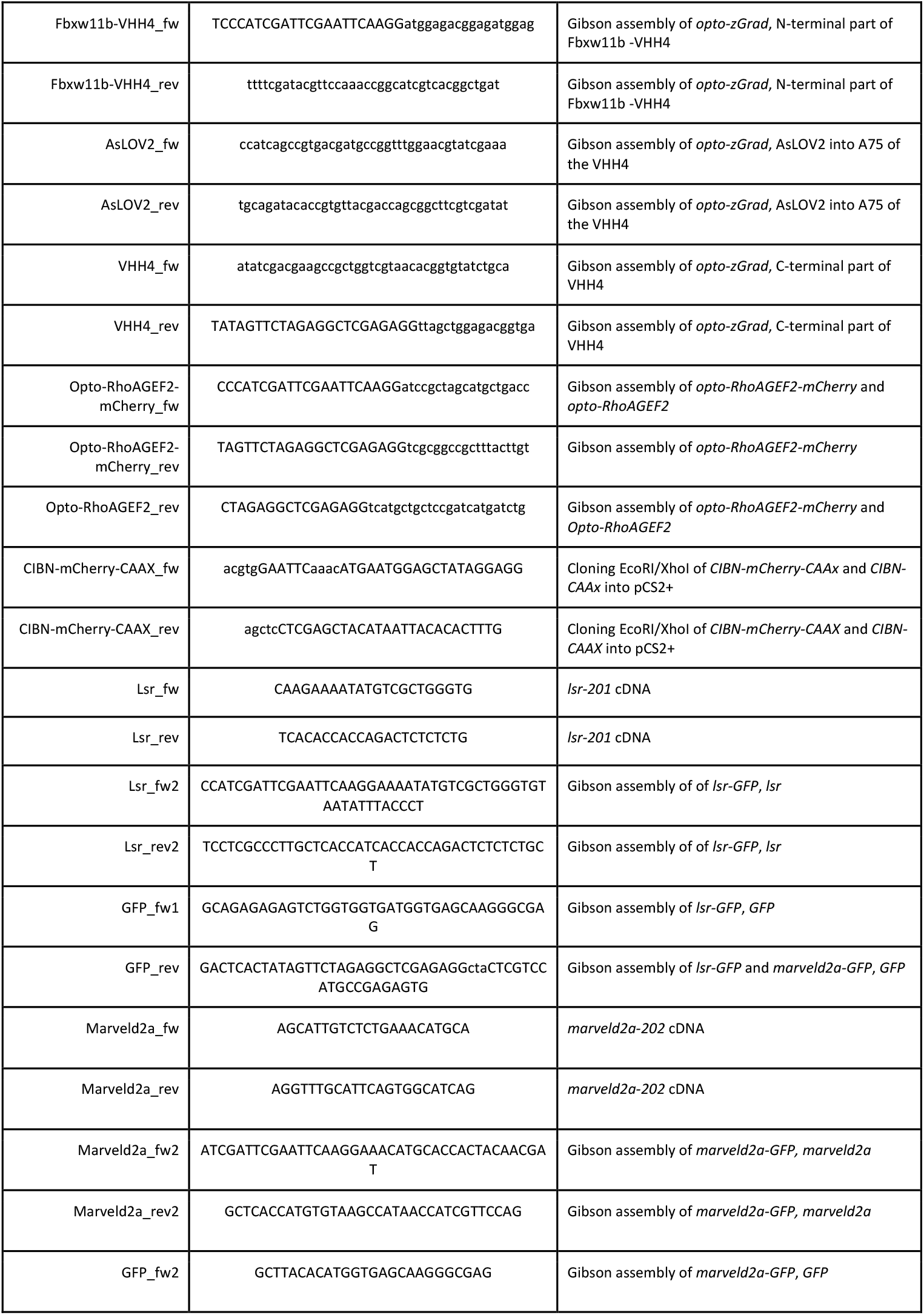

### Embryo microinjections and explant formation

Zebrafish embryos were injected using glass capillary needles (30-0020, Harvard Apparatus, MA, USA) that were pulled with a P-97 needle puller (Sutter Instrument) and attached to a PV820 microinjector system (World Precision Instruments). Microinjections of mRNAs were performed at the one-cell stage into the yolk. mRNAs were synthesized from linearized plasmids using the mMESSAGE mMACHINE™ SP6 Transcription Kit (Invitrogen, Cat. No. AM1340). The following mRNAs were injected: i) Cell membrane labeling: 50–70 pg *membrane-RFP* ^82^, *membrane-GFP* ^83^; ii) Nuclear labeling: 20–40 pg *H2A-BFP* ^84^, *H2A-mCherry* ^85^, or *H2B-GFP* ^86^; iii) Tricellular junction labeling: 17.5 pg *lsr-GFP* and 17.5 pg *marveld2a-GFP*; iv) *α*-catenin-citrine degradation: *Gt(ctnna-citrine)*^*ct3a*^ homozygous embryos were microinjected with 35 pg *opto-zGrad* and co-injected with a nuclear or cell membrane marker to ensure injection quality; v) RhoGEF activation to decrease relative surface tension: 50– 70 pg *CIBN-CAAX* or *CIBN-mCherry-CAAX* co-injected with 17.5 pg *opto-RhoGEF-CRY2*, or without it as a control. To label the interstitial fluid, high-stage dechorionated embryos were microinjected with 0.5 nL of 0.6 µg/µL Dextran-Alexa Fluor® 647 (10,000 MW; Invitrogen, Cat. No. D22914) into the blastoderm interstitial fluid space. Explants were prepared by isolating the blastoderm from high-stage embryos and maintained in 1x Danieau’s medium. Staging was determined relative to their sibling control embryos. Interstitial fluid labelling of the explants was performed by microinjection as described above, 30 min after blastoderm excision to allow tissue rounding up and would healing.

### Pharmacological treatments

To increase interstitial fluid fraction, high-stage dechorionated embryos were injected into the blastoderm interstitial space with 0.5 nL of 400 mM D-Mannitol (Sigma-Aldrich, M4125-100G) supplemented with 0.6 µg/µL of Dextran-Alexa Fluor® 647 to label the interstitial fluid. To decrease interstitial fluid, dechorionated embryos at the 2-cell stage were incubated with 1 mM Ouabain (Sigma-Aldrich, 4995-1GM) until sphere stage, followed by interstitial fluid labelling (as described above).

### MPA and viscosity measurements

Blastoderm viscosity was measured by micropipette aspiration based on previously established protocols ^5,19,24^. Briefly, embryos were placed on 3% methylcellulose-coated glass coverslips in 1x Danieau’s solution on an inverted Leica SP5 microscope. A fire-polished, passivated (with heat inactivated FBS) micropipette of 35 mm inner diameter, 30 bent, with a spike end (Biomedical Instruments) was inserted into the blastoderm just below the EVL. The micropipette movements were controlled by motorized micromanipulators (Eppendorf Transferman, Nk2). Upon insertion of the pipette in the blastoderm, an aspiration pressure of 150 Pa was immediately applied using a Microfluidic Flow Control System Pump (Fluigent, Fluiwell) (with negative pressure ranging from 7-750 Pa, a pressure accuracy of 7 Pa and change rate of 200 Pas_-1_) and the Dikeria micromanipulation software. The value of the applied pressure (150 Pa) was set according to prior test aspiration experiments, where the applied pressure was continuously increased in a stepwise process (10 Pa / 20 s) until the aspirated tissue started flowing into the pipette. Pressure was applied until the tissue flowed into the pipette at a constant velocity (for 3 min, except the cases of very fast deformation) and then pressure was immediately released. Images for monitoring the aspiration and relaxation of the tissue were acquired every 500 ms. Viscosity calculations were performed as previously described ^5,19,24^, using a customized Fiji macro to calculate changes in tongue length during aspiration and relaxation over time. The slope of the aspiration curve (*L*_*asp*_) at the point of constant flow depends on the viscosity η, *L*_*asp*_ = *R*_*p*_(Δ*P* − *P*_*c*_)/3πη, with *R*_*p*_ the radius of the pipette, Δ*P* the applied pressure and *P*_*c*_ the critical pressure. During the relaxation, the tissue retracts at a velocity *L*_*ret*_ = *R*_*p*_(*P*_*c*_)/3πη. From the aspiration and retraction rates, viscosity can be calculated as η = *R*_*p*_Δ*P*/3π(*L*_*asp*_ + *L*_*ret*_). The data plotted in Fig. 1L were reused from ^19^.

### Immunostaining

Samples were fixed in Glyoxal solution mix (GS). For a 4 mL fixative preparation, 2.835 mL H_2_O, 0.789 mL 100% EtOH, 0.313 mL Glyoxal (40% stock solution; Sigma-Aldrich, 128465), and 30 µL Acetic Acid (100%) were mixed; pH was finally adjusted to 4–5 with 1 M NaOH based on ^87^. Dechorionated zebrafish embryos and explants were incubated in GS for 1 h (30 min on ice and 30 min at room temperature), followed by a 15 min quenching step in 100 mM NH_4_Cl / 100 mM Glycine. Explants were pre-permeabilized for 2 h in 1× PBS / 1% Triton X-100 / 2% BSA, followed by overnight incubation in blocking/permeabilizing solution (B/PS; 1× PBS / 0.5% Triton X-100 / 2% BSA) for both explants and embryos. Primary antibodies were incubated overnight at 4°C in B/PS. The following primary antibodies were used for this study: Mouse anti-E-Cadherin (BD Biosciences, Cat. No. 610181, 1:200), Mouse anti-ZO-1 (Thermo Fisher Scientific, 33-9100, 1:100), Rabbit anti-Phospho-Myosin Light Chain 2 (Ser19) (Cell Signaling, 3671, 1:100), and Rabbit anti-Pan-Cadherin (Sigma-Aldrich, C3678, 1:200). After three washes in B/PS (30 min each), samples were incubated at 4°C with secondary antibodies in B/PS overnight. The following secondary antibodies were used: Goat Alexa Fluor 546 anti-rabbit (Thermo Fisher Scientific, A11010, 1:500), Goat Alexa Fluor 488 anti-rabbit (Thermo Fisher Scientific, A11008, 1:500), Goat Alexa Fluor 546 anti-mouse (Thermo Fisher Scientific, A11003, 1:500), Goat Alexa Fluor 488 anti-mouse (Thermo Fisher Scientific, A11001, 1:500). Finally, samples were washed in 1x PBS / 0.5% Triton X-100, followed by a 30 min incubation in 300 nM DAPI / 1x PBS staining solution (Invitrogen, D1306) in PBS with 1x Rhodamine-Phalloidin (Abcam, ab235138) if required. After a final wash with PBS, samples were mounted in 80% glycerol / 0.4% N-propyl-gallate / 1× PBS. All steps were performed with gentle horizontal rocking.

### Image acquisition

Dechorionated embryos and explants were embedded in 0.6% low melting point agarose (Invitrogen, Cat. No. 16,520-050) on a customized agarose mold in a 60 mm dish (Greiner, 628102). Mounted embryos and explants were kept in an incubation chamber at 28.5°C throughout acquisition. Live imaging was performed on an upright ZEISS LSM 980 with Airyscan 2 with Axio Examiner. Confocal live imaging of deep cells to reconstruct connectivity maps was performed with an W Plan-Apochromat 20x/1.0 Corr DIC M27 75mm objective. Immunostaining samples were imaged with a LD LCI Plan-Apochromat 40x/1,2 Imm Korr DIC M27 Zeiss objective both in Confocal and Fast Airyscan super-resolution modules. Contact length fluctuations were imaged with an inverted ZEISS LSM 980 with Airyscan 2 (MPLX SR-8Y mode) and C-Apochromat 40x/1.2 W autocorr FCS M27 objective or with an upright ZEISS LSM 980 with Airyscan 2 (MPLX SR-4Y mode) and W Plan-Apochromat 20x/1.0 Corr DIC M27 75mm objective. Timelapses were acquired over at least 10 min with an interval of 10 sec. Images were acquired in ZEN3.3 (blue edition) software (Carl Zeiss) and processed using Fiji (https://fiji.sc/).

### Optogenetics

Embryos were kept in the dark until the selected developmental stage. To prevent photoactivation, all sample handling and mounting was performed under red light filters (Lee Filter 106, Primary Red) to block full bright-field illumination. Pre-imaging photoactivation was performed at sphere stage in most of the experiments, using a 50 W/60 Hz blue LED lamp in the incubator. For imaging, embryos were oriented for transverse and sagittal blastoderm views according to the experiment and imaged/photoactivated from sphere to shield stage. For *α*-catenin-citrine degradation, *Gt(ctnna-citrine)*^*ct3a*^ homozygous embryos were microinjected with opto-zGrad as described above. Opto-zGrad photoactivation and imaging of *α*-catenin-citrine degradation were carried out with 488 nm light pulses (laser power set between 6–8%, corresponding to an out-of-objective power of 104–140 µW) every ~3 min, with a stack size of ~45 µm (4 µm spacing). Dark effects were sometimes observed in a concentration-dependent manner. For opto-RhoGEF experiments, *Gt(ctnna-citrine)*^*ct3a*^ embryos microinjected with the opto-RhoGEF system were photoactivated/acquired with a 5-8% 488 nm laser (corresponding to 86-140 µW) in a total z stack of 45 µm (z-stacks (4 µm spacing), with pulses every ~5 min. Concomitant with photoactivation, 647 nm, 561 nm, and 405 nm excitation was used to image Dextran-Alexa647 and membrane-RFP and/or Histone-BFP when present.

### Data analysis and quantification

All acquired data were processed using Fiji ^88^ and Cellpose v2 and v3 ^89,90^. R (version 4.2.2) was used for data analysis and plotting (ggplot2 version 3.5.1) ^91^.

#### Image segmentation

Blastoderm images were segmented using Cellpose v2 and v3, with manual corrections being performed where necessary.

#### Reconstruction of connectivity maps and rigidity analysis

Cell connectivity was defined on 2D confocal sections of the 2^nd^-3^rd^ deep-cell layer, in which the nuclei, cell–cell contacts and interstitial fluid accumulations were differentially labelled. The image from the interstitial fluid channel was converted to a binary image and, if necessary, processed by a median filter of 2-pixel width in Fiji to emphasize the interstitial fluid gaps. Cells within the same focal plane that had no interstitial fluid between them were considered as contacting cells. For small contact areas, the non-binarized image was compared for confirmation. The connectivity networks were reconstructed based on the above described Cellpose segmentation. Adjacency between labels was assessed using a custom-made plugin in Fiji that uses the MorpholibJ function “Region Adjacency Graph” ^92^. The plugin allowed labels to be manually added/removed and the single links to be manually curated to achieve corrected networks. The plugin generates a text file with cell centroids (x,y) for each cell label and a text file for contact adjacency (based on cell labels) used later for rigidity percolation analysis. The coordinates of each cell centroid and their contacting neighbors were identified and plotted using the standard python package matplotlib.

#### Cell connectivity and tissue rigidity

Cell connectivity was calculated in each confocal section as the total number of contacts (defined as described in connectivity map reconstruction) divided by the total number of cells in the image. Normalized connectivity (*< k >*) was calculated in each confocal section as connectivity C divided by the maximum potential connectivity (*k*_*max*_) (computed as described in ^19^). The rigidity analysis to identify floppy and rigid areas, Giant Rigid cluster, and cluster size distributions was performed on the connectivity maps using a python version of the pebble algorithm (pebble.py).

#### Cell fraction, tissue porosity and lumen formation

Cell fraction *ϕ* was calculated as: *ϕ* = 1 − *ff*, where *ff* is fluid fraction. *ff* was calculated from 2D confocal sections at the 2^nd^ deep cell layer, where the interstitial fluid channel (labeled with Alexa Fluor 647 Dextran) was first converted into a binary image as described above. A signal intensity histogram analysis of the whole binary image was performed in Fiji, and the *ff* was obtained by the average gray value of the histogram normalized to the maximum signal intensity of the histogram. For simulated tissues (Fig. 1G), we created a .ps figure from Surface Evolver tiling and segmented the image to identify cells and fluid. The fluid fraction was then computed as the ratio of fluid-labeled pixels to the total number of pixels in the image. Tissue porosity was quantified as a function of the number of pockets per cell times the mean cell perimeter fraction in contact with interstitial fluid. Lumen size was quantified upon binarization of the interstitial fluid as described above, followed by a connected regions analysis in Fiji to identify the largest region composed of interstitial fluid signal.

#### Relative surface tension

The cell-cell contact angle was calculated in degrees from two independent 2D confocal sections using the angle tool in Fiji, as indicated in Figure 1E, and then converted to radians. The relative surface tension was then calculated following the methodology developed in ^36^ as: *α* = cos (θ/2).

#### Morpho-feature quantification

Cell and tissue morphological features were extracted from segmented microscopy images using a custom Python pipeline designed for high-throughput feature extraction. The analysis was performed using imageio (2.34.2), scikit-image (0.24.0), numpy (1.26.4), pandas (2.2.2), and scipy (1.14.0) in Python 3.12.4. Labeled segmentation masks generated by Cellpose were upscaled 4×4 to avoid loss of small interstitial fluid pockets and processed iteratively using os. Each image was loaded with imageio.imread(), and individual cell labels were extracted using numpy.unique(). Cellular morphological properties - including area, perimeter, orientation, major axis length, minor axis length, and circularity - were computed using skimage.measure.regionprops. To detect tricellular junctions, a junction map was generated using scipy.ndimage.generic_filter, identifying locations where three or more cells meet. Subsequently, these values were normalized by the total number of triangles in the network, obtained from the connectivity map described above and the mathematical derivation presented in the Supplementary Theory Note. To analyse cell-cell and cell-fluid contacts, the background (interstitial fluid) of each image was segmented using skimage.measure.label(). Cell labels were dilated by one pixel, and the overlap with neighbouring cells and background regions was calculated. This enabled quantification of cell-cell contact proportions and identification of the corresponding background regions. The same approach was used to analyse fluid pockets. For each input segmentation, the extracted parameters were summarized as mean and standard error of the mean (SEM). The values for area, perimeter, circularity and adjacent labels were extracted for individual cells (labels) and fluid pockets. During processing of the data, the following parameters were normalized: number of fluid pockets (divided by the number of cells), pocket size (pocket area divided by the fluid area), percentage of closed triangles (as described above), and the number of tricellular junctions per cell (total number of tricellular junctions divided by the number of cells). The mean cell aspect ratio was calculated as the ratio of the mean major axis to the mean minor axis length.

#### Dimensionality reduction and clustering

Principal component analysis (PCA) was performed on various metrics describing qualitatively various aspects of cell and tissue morphology using the prcomp() function of the Stats package in R (R version 4.2.2 for all analyses), with centering and scaling to unit variance for normalization. Metrics acting as control parameters of material state phase transitions (relative surface tension, cell fraction, cell shape index, connectivity) were not included. Metrics included in the PCA were: normalized number of TCJs, percentage of closed triangles, mean cell perimeter fraction in cell-cell contacts, fraction of cells in contact with fluid (thresholded), cell circularity, maximum fluid pocket size (normalized), number of pockets per cell, and cell aspect ratio. PCA coordinates were later merged with the raw dataset to allow color-coding and filtering for specific conditions and parameters. The tissues were represented in a 2D morphospace by plotting them against PC1 and PC2 using ggplot2. Uniform Manifold Approximation and Projection (UMAP) was performed on the same scaled metrics using the umap package (version 0.2.10.0) in R, with n_neighbors set to 8 and min_dist to 0.1. UMAP coordinates were later merged with the raw dataset to allow color-coding and filtering for specific conditions and parameters. The tissues were represented in a 2D morphospace by plotting them against UMAP1 and UMAP2 using ggplot2. To assign each tissue to a cluster, k-means clustering was performed on the UMAP1 and UMAP2 coordinates using the kmeans() function of the stats package in R, specifying 4 cluster centres based on the number of material phase space regimes.

#### Cell-cell contact length fluctuations

Cell-cell contact length fluctuations were calculated from 2D confocal time series acquired using the Fast Airyscan mode (10 s interval, 10 min time lapse duration). For each time point, the contact length was measured using the free-hand line tool in Fiji (as shown in Figure 2H, 3E). Then, contact lengths were normalized to the average of the contact lengths during the 10 min duration and the coefficient of variation (CV) for each contact during the 10 min measurement was calculated. The average CV of at least 12 contacts from at least 3 embryos per time point and condition was plotted (Figures 2I, J, 3F, S3F).

#### *α*-catenin quantification

To quantify *α*-catenin degradation with opto-zGrad and assess photobleaching effects under our imaging settings, *α*-catenin-citrine intensity was measured every ~10 min as the mean grey value within a defined region of interest on the sum projection of four consecutive z-layers (16 µm), starting from the second layer of cells beneath the most apical region of the embryo, using Fiji. For each embryo, the values were normalized by the value of their respective starting imaging intensity.

### Statistics and reproducibility

The statistical analyses were performed with GraphPad Prism 10.0 and R (version 4.2.2) with the following packages: ggplot2 3.5.1, ggpubr 0.6.0, broom 1.0.7, car 3.1-3, FSA 0.9.6, multcomp 1.4-28, DescTools 0.99.54, dplyr 1.1.4, and stats 4.2.2. Statistical details of experiments are reported in the figures and figure legends. Sample sizes are provided in the figure legends, and no statistical test was used to determine sample size. The biological replicate is defined as the number of embryos, as stated in the figure legends. No inclusion or exclusion criteria, randomization, or blind allocations were applied, and all analyzed samples were included. Unless differently stated in the figure legends, the graphs show mean ± SEM, and the error bars are calculated and shown based on the number of cells or embryos, as indicated. The statistical test used to assess significance is stated in the figure legends and was chosen after testing each group for normality using the Shapiro-Wilk test and for homogeneity of variances using Levene’s test. If any condition had n ≤ 3, Levene’s test was omitted, and a non-parametric test was used. For comparisons between two groups, a two-tailed Student’s t-test was used for parametric distributions with equal variances, a Welch’s t-test for parametric distributions with unequal variances, and a Mann–Whitney U-test for non-parametric distributions. For multiple pairwise comparisons, an ANOVA followed by Dunnett’s test was used for parametric distributions, while a Kruskal-Wallis test followed by Dunn’s test with Dunnett’s adjustment or Bonferroni correction (for pairwise comparisons) was applied for non-parametric distributions. For correlation analysis, Spearman’s correlation was performed. At least three independent experiments were conducted unless stated differently in the figure legend. This information is also stated in the figure legends.

### Reagents, data and code availability

All reagents, codes and data generated in this study will be publicly available upon manuscript publication.

## SUPPLEMENTARY FIGURES AND FIGURE LEGENDS

**Figure S1:**
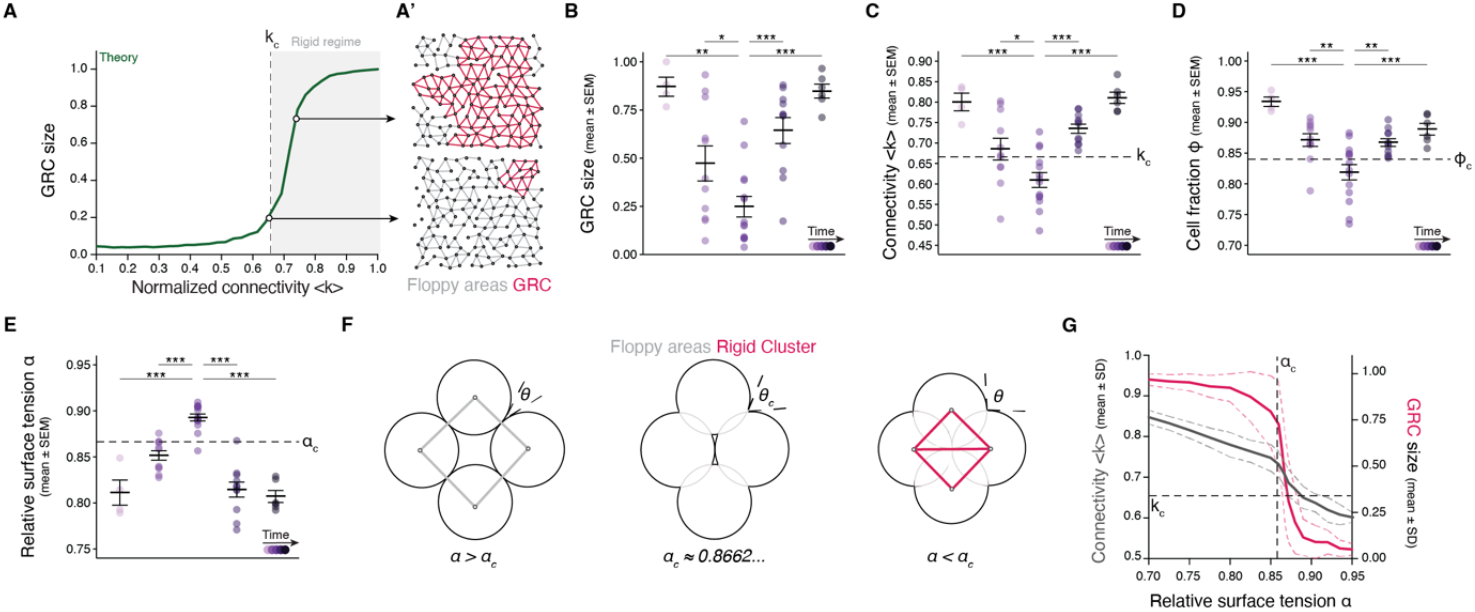
Quantitative analysis and theoretical relationships between multiple control parameters and tissue rigidity and viscosity. **(A)** Plot of the fraction of the network occupied by GRC as a function of normalized connectivity <*k*> in simulated random 2D triangular lattices like the exemplary networks shown in A’. The gray-shaded area indicates the network rigid regime above the critical connectivity point (dashed line). **(B-E)** Plots of experimental data from high-until shield-stage embryos quantified for (B) the fraction of the network occupied by GRC, (C) normalized connectivity < *k* >, (D) cell fraction *ϕ*, and (E) relative surface tension *α*. Embryo numbers as in Fig. 1. **(F)** Numerical simulations for the morphology of a 4-disk rhombus cluster at different relative surface tension *α* values. By decreasing *α* below *α*_*c*_, a new contact is formed, rigidifying the small cluster. **(G)** Plot of normalized connectivity <*k*> (left Y-axis, gray) and fraction of the network occupied by GRC (right Y-axis, red) as a function of relative surface tension *α* for simulated cell arrays like those depicted in Fig. 1F, showing a sudden increase in connectivity and GRC size at *α*_*c*_. Floppy areas are illustrated in gray and GRC in red. Kruskal-Wallis with Dunn’s post hoc test (Dunnett adjustment) (B); ANOVA with Dunnett’s post hoc test (C-E).

**Figure S2:**
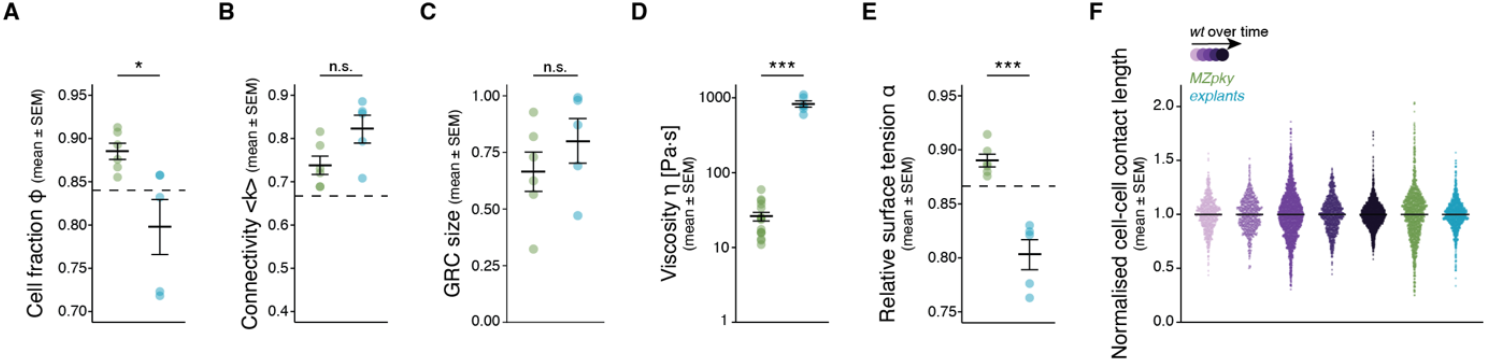
Quantitative analysis of cell and tissue biophysical properties during the rigidity transition. **(A-F)** Plots of (A) cell fraction *ϕ*, (B) normalized connectivity <*k*>, (C) fraction of the network occupied by the GRC, (D) blastoderm viscosity, (E) relative surface tension *α*, and (F) cell-cell contact lengths for *MZpky* and explants. Embryo and contact numbers as in Fig. 2. Dashed lines: critical points. Mann-Whitney U-test (A); Student’s t-test (B, C, E); Welch’s t-test (D).

**Figure S3:**
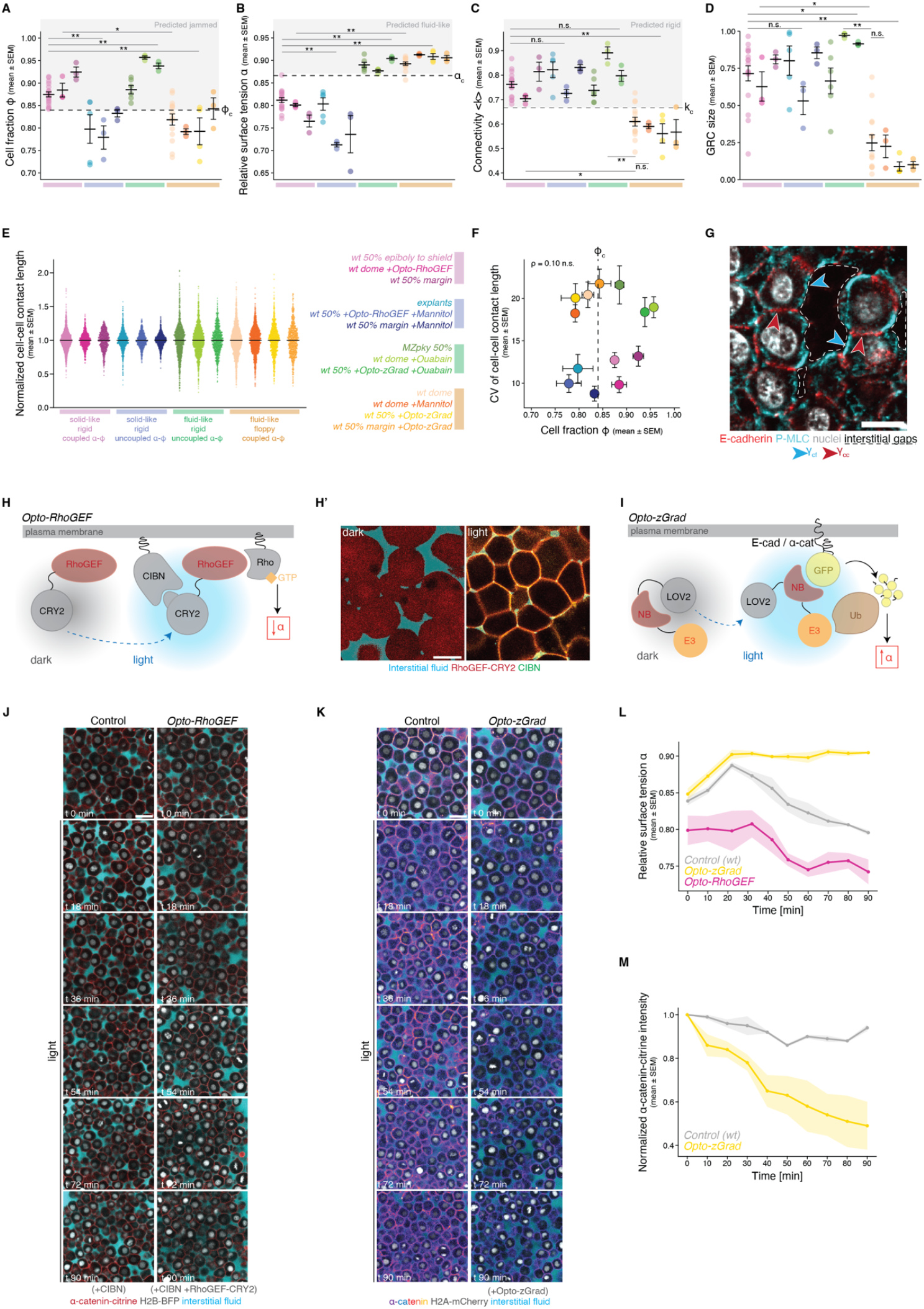
Quantification of control and order parameters in all experimental conditions and characterization of the optogenetic tools. **(A-E)** Plots of quantifications of cell fraction *ϕ* (A), relative surface tension *α* (B), normalized connectivity <*k*> (C), fraction of the network occupied by the GRC (D), and normalized cell-cell contact lengths (E) for all experimental conditions listed (embryo and contact numbers as in Fig. 3). **(F)** Plots of the coefficient of variation (CV) in cell-cell contact length for the conditions shown in (E) as a function of cell fraction *ϕ* (embryo and contact numbers as in Fig. 3). **(G)** Exemplary high-resolution 2D confocal section of a *wt* embryo stained for E-cadherin and phosphorylated myosin (P-MLC) showing differential localization at the cell-cell and cell-interstitial gap interfaces. The arrows indicate where the cell-cell tension γ_cc_ (red) and the cell-fluid tension γ_cf_ (blue) act. **(H)** Schematic diagram of the opto-RhoGEF tool, which combines a membrane-tagged CIBN and RhoGEF-CRY2 ^48^. **(H’)** Upon light illumination, CIBN binds to CRY2, translocating RhoGEF to the membrane. **(I)** Schematic diagram of the opto-zGrad tool, which is composed of an AsLOV2 (LOV2) domain attached to a GFP nanobody (NB) and a domain for proteasomal degradation targeting (E3). **(J)** Exemplary 2D confocal sections over the course of 90 min of illumination, from sphere to 30-50% epiboly, for both control (*wt* + CIBN) and opto-RhoGEF (*wt* + CIBN + RhoGEF-CRY2). Fluidization occurs at t = 18 min. Interstitial fluid labeled with dextran-647. **(K)** Exemplary 2D confocal sections of blastoderms from zebrafish embryos from a knock-in line for *α*-catenin-citrine, either without (control) or with expression of opto-zGrad. The 90 min timelapse of the illumination corresponds to the stages of sphere until 30-50% epiboly, with doming at t = 18 min. Membranes labeled with *α*-catenin-citrine, interstitial fluid with dextran-647. **(L)** Plot of the quantification of relative surface tension *α* over time (n = 140-210 angles per time point; embryo numbers: N = 4 control, N = 3 opto-zGrad, N = 3 opto-RhoGEF). **(M)** Plot of the quantification of *α*-catenin-citrine fluorescence intensity in control and opto-zGrad embryos over time (N = 3 embryos per condition). Mann-Whitney U-test (A-D); ρ: Spearman correlation (F). Scale bars: 10 µm (G), 20 µm (H’) 30 µm (J, K).

**Figure S4:**
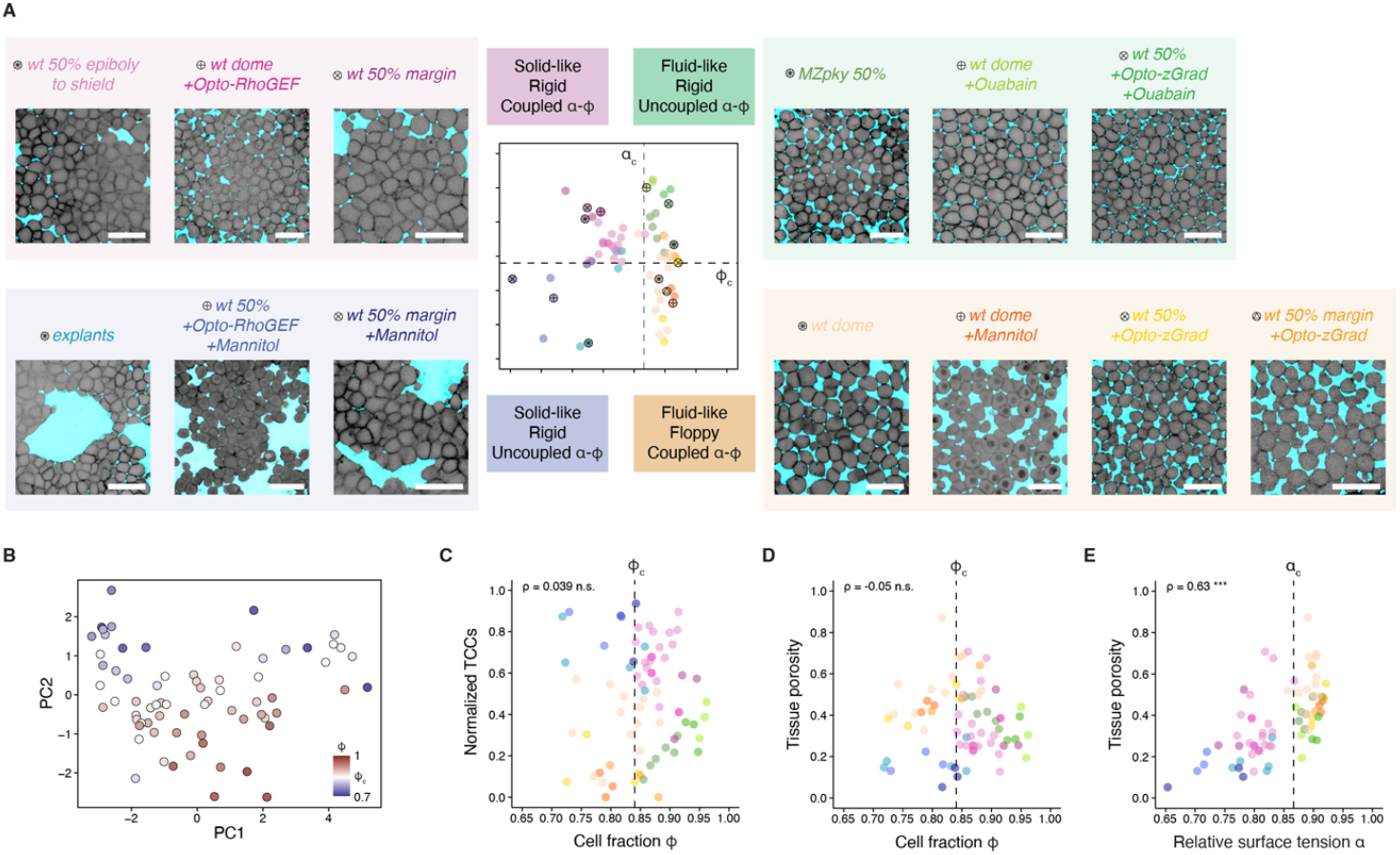
Phase transition-dependent discretization of tissue morphology and organization. **(A)** Exemplary 2D confocal sections showcasing tissue morphology for the experimental conditions occupying each material regime in the phase diagram (embryo numbers as in Fig. 3). Interstitial fluid labeled with dextran-647, membranes with *α*-catenin-citrine, membrane-GFP, or Lifeact-eGFP. **(B)** Morphospace of all tissues shown in the material phase diagram in (A) plotted against PC1 and PC2, color-coded by cell fraction *ϕ*. **(C)** Plot of the fraction of TCCs that are closed as a function of cell fraction *ϕ* for all the tissues shown in (A). (**D-E**) Plots of tissue porosity as a function of cell fraction *ϕ* (D) and relative surface tension *α* (E). Dashed lines: critical points. ρ: Spearman correlation (C-E). Scale bars: 50 µm.

**Figure S5:**
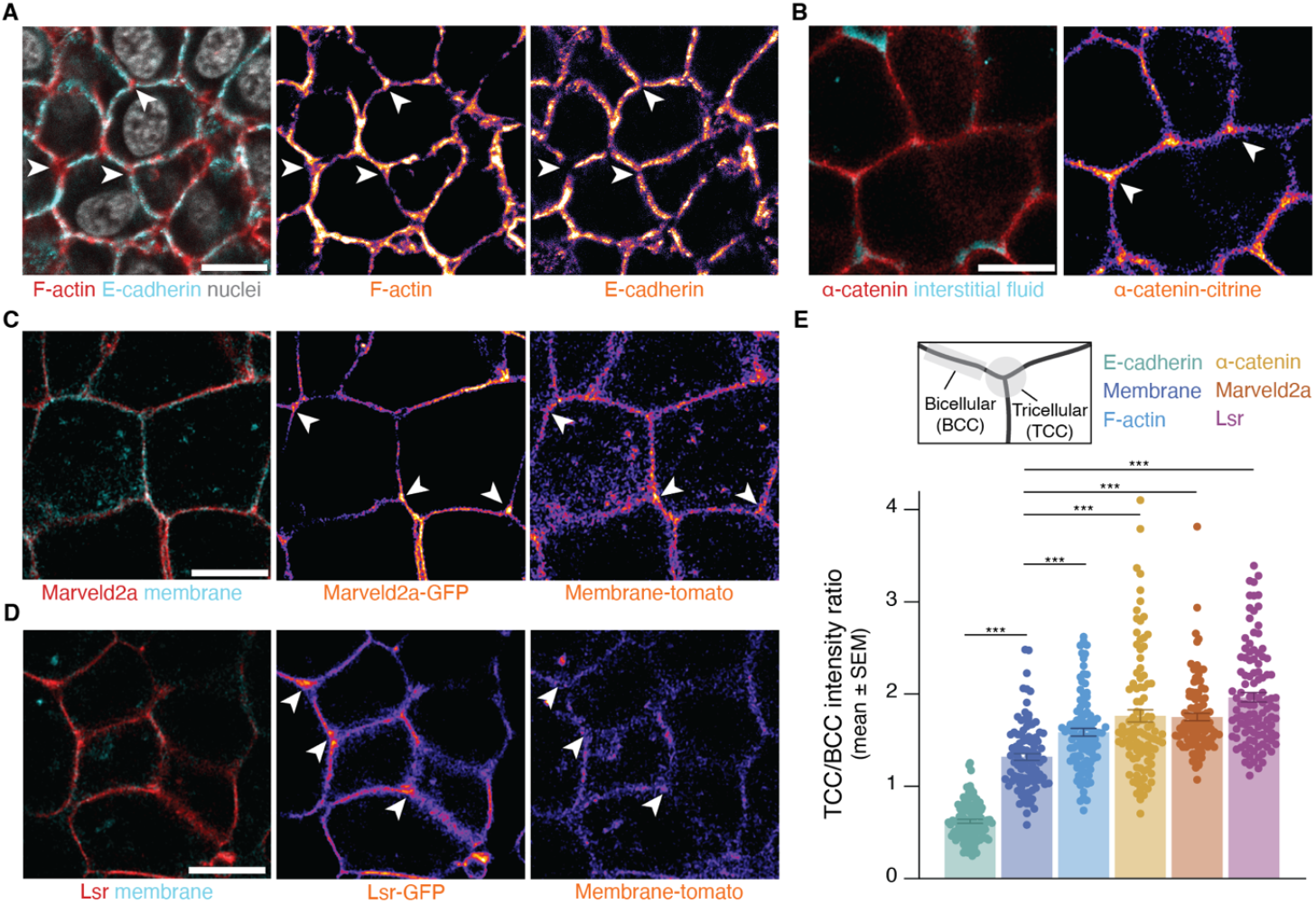
Accumulation of tricellular junction-specific proteins at the tricellular contacts of pluripotent cells undergoing an adhesion-driven tissue rigidification. **(A-D)** Exemplary high resolution 2D confocal sections of *wt* shield-stage embryos (A) stained for F-actin (phalloidin) and E-cadherin, or live imaged and labeled for (B) *α*-catenin and interstitial fluid, (C) Marveld2a and membrane, and (D) Lsr and membrane. White arrowheads indicate tricellular junctions. **(E)** Plot of the quantification of fluorescent protein intensity at the tricellular (TCC) vs. the bicellular (BCC) contacts for the proteins shown in (A-D) (n = 50 cells per condition, N = 4 embryos each). Scale bars: 10 µm (A-D). Kruskal-Wallis test (E).

## I. CELL-CELL CONTACT ENERGY

We study non-confluent tissues, where cells are in contact among them and with the interstitial fluid. We are interested in cell-fluid interface and cell-cell contacts. To that end, we assume that energy of the system can be approximated by the Hamiltonian of a soap bubble [1, 2]:

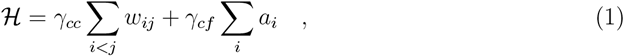

being the first sum performed over all cell-cell potential contacts, w_*ij*_ the contact area between cells i and j and a_*i*_ the area of cell i exposed to the interstitial fluid. Throughout the paper we consider that cell contacts are governed by Young-Dupré’s law. This law states that, in a force-balance situation, the contact angle between the membranes of two cells in contact, θ, and the relation between the strength of the cell-cell surface tension, γ_*cc*_, and cell-fluid surface tension, γ_*cf*_, are related through [2–4]:

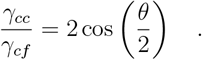

**Suppl. Note Fig. 1:**
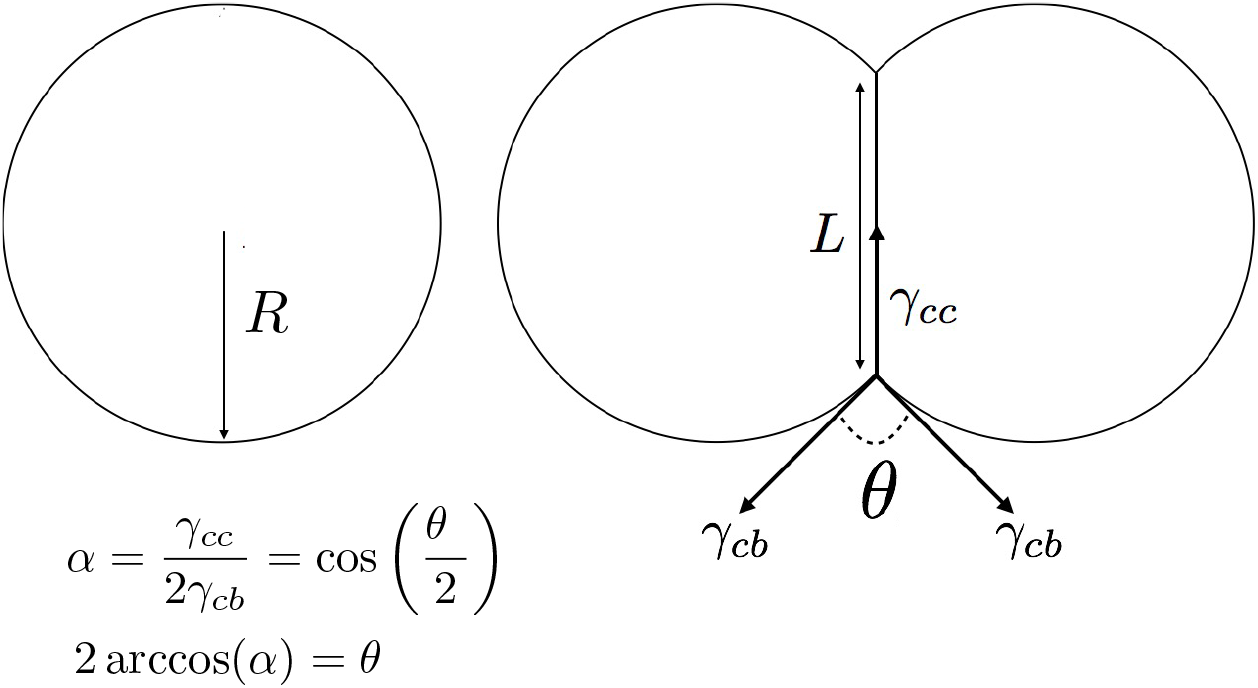
Schematic representation of the Young Dupré relation.

In Suppl. Note Fig. (1) we detail schematically each of these terms. From the above relation, one can derive a non-dimensional parameter, *α*, describing the relative strength of the cell-cell surface tension, γ_*cc*_ and cell-fluid surface tension, γ_*cf*_ :

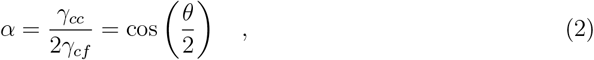

and, consequently, re-write the Hamiltonian in a non-dimensional form:

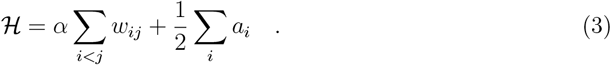

Key to all this framework is that the non-dimensional parameter *α* can be inferred from the cell-cell contact angles, an observable that can be empirically extracted. This enables us to perform predictions on structural properties of the arrangements of cells in contact only considering geometric grounds. Such derivations are valid in a context with no net stresses applied over the tissue.

## II. ADHESION-INDUCED RIGIDIFICATION

### A. Generic rigidity

We will use the concept of *generic rigidity*, a graph-theoretic concept [5]. Suppose a network G^′^(V ^′^, E^′^) made of a set V ^′^ of nodes and a set E^′^ of links between nodes. A *spanning subgraph* is a subgraph G(V, E) of G^′^(V ^′^, E^′^) by which V = V ^′^ and E^′^ ⊆ E. An *induced subgraph* g(V_*g*_, E_*g*_) of G^′^(V ^′^, E^′^) is a subgraph by which V_*g*_ ⊆ V and in which all the links between elements of V_*g*_ existing in G^′^ are present. Finally, a graph is *connected* if there is a path between any pair of nodes of V ^′^.

As we work using 2D projections, we ground our reasoning on the plane. Consider that links impose spatial constraints. In d dimensions, two nodes v_*i*_, v_*j*_ ∈ E will have a priori d spatial degrees of freedom each, but if {v_*i*_, v_*j*_} ∈ E, that is, there is a link between v_*i*_ and v_*j*_, then, the degrees fo freedom of v_*i*_ and v_*j*_ will be at most d − 1, because the link acts as spatial constraint for the two nodes. A network is called generically rigid if the set of links absorbs all the degrees of freedom of the nodes in d dimensions, such that nodes do not have independent movements. This implies that, if nodes are interpreted as rigid bars or springs at rest, no deformations of the structure are possible at no energy cost; i.e., the Young modulus of the whole network is > 0 [6]. In general, a network G^′^(V, E^′^) embedded in a 2-dimensional space will be generically rigid if there is a connected, spanning subgraph G(V ^′^, E), with E ⊆ E^′^, by which:

1. |E| = 2|V ^′^| − 3
2. For every subset V_*g*_ ∈ V ^′^, with |V_*g*_| ≥ 2, the induced subgraph g (with link set E_*g*_ ∈ E^′^) is such that |E_*g*_| ≤ 2|V_*g*_| − 3.

This theorem is due to Laman [7]. We emphasize that generic rigidity is a purely topological property of the network.

With these tools in hand, we will look at a specific graph, a rhombus –see Suppl. Note Fig. (2A). In this case, it is easy to check that the rhombus is not generically rigid because even if we consider the whole graph, then: |E^′^| = 4, |V ^′^| = 4, and, thus, |E^′^| < 2|V ^′^| − 3, forbidding the existence of a spanning subgraph satisfying the first condition of Laman’s theorem. In the case of the graph of Suppl. Note Fig. (2B,C), with an accessory link connecting the central nodes, |E^′^| = 5, and the whole graph itself satisfies the first condition of Laman’s theorem. A simple counting argument throughout all the potential configurations can be used to check that any induced subgraph satisfies the second condition of Laman’s theorem. Therefore, the graph in A/ is floppy and the ones in B/ and C/ are generically rigid.

### B. Generic rigidity arising from cell-cell adhesion changes

We want to explore how the reduction in *α* may affect the topology of cell-cell contacts and, in consequence, the potential rigidity properties. To that end, we use the toy motif of 4 cells in a 2D projection that is analytically tractable. This will allow us to determine a critical point in *α*. Further, using numerical simulations, we will see that the result scales to arbitrary cell arrangements.

Let us suppose that we have a floppy motif made of 4 cells –see see Suppl. Note Fig. (2D). The network of cell contacts corresponds to the one shown in see Suppl. Note Fig.(2A), for which we have shown it is not generically rigid. We want to know at what *α* the motif rigidifies. That is: at which *α* a new contact is created such that its cell-cell contact network corresponds to the one shown in see Suppl. Note Fig. (2B,C). To that end, we start with a symmetric, rhombus-like configuration of 4 disk shaped cells with only pointwise contacts –see Suppl. Note Fig. (2A,D)– of radius 1 located at 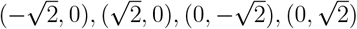, respectively. To mimic the effect of the increase of adhesion, we will progressively bring the disks located left and right of the vertical axis closer and closer, allowing overlap. The new configuration of the centers of the cells will be 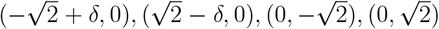, respectively, and we keep the radius invariant. The bisectrix line between two overlapping circles will represent the cell-cell contact region without any loss of generality, as we are only interested in the angle between the two circles and, therefore, conservation of area is not required, although it can be achieved by just applying a rescaling, which would not affect the configurations of angles. The disks on top and at the bottom of the rhombus remain fixed. Eventually a new contact is created, when the location of the centers of mass is 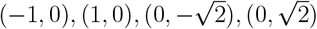, leading |E| = 5 and, therefore, according to the derivation provided in the previous section, (generically) rigidifying the motif –see Suppl. Note Fig. (2B,E,F). Now it remains to compute the angle between circles that results from the obtained configuration. From that, we can straightforwardly, applying Young-Dupre’s law, compute *α*_*c*_. The critical point at which the new contact is corresponds to:

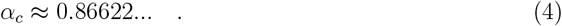

**Suppl. Note Fig. 2:**
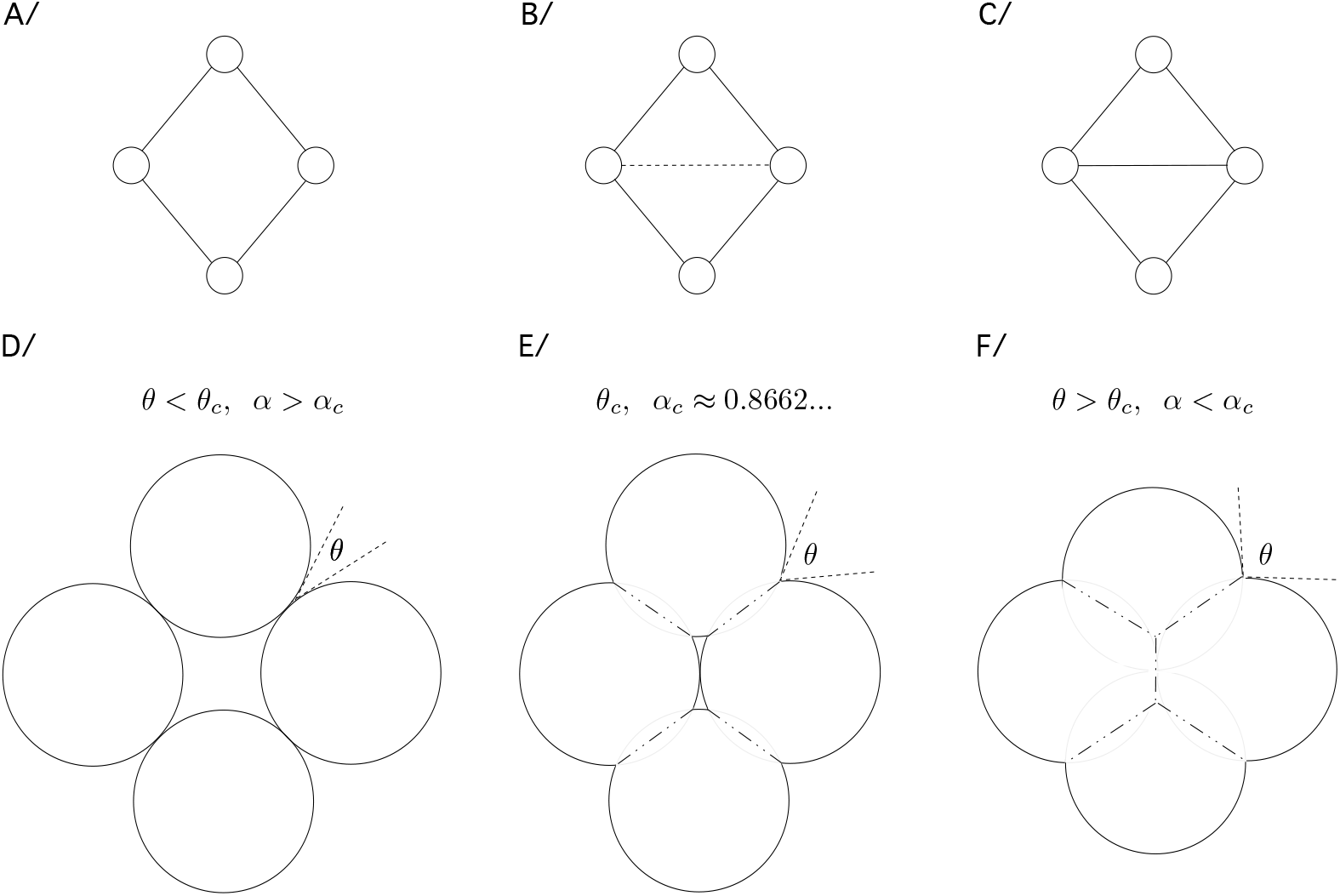
Topological and geometric effects of the decrease of *α*.

The whole computation is based on finding the θ_*c*_ using standard geometric considerations. In Suppl. Note Fig.(3) we detail the reasoning.

**Suppl. Note Fig. 3:**
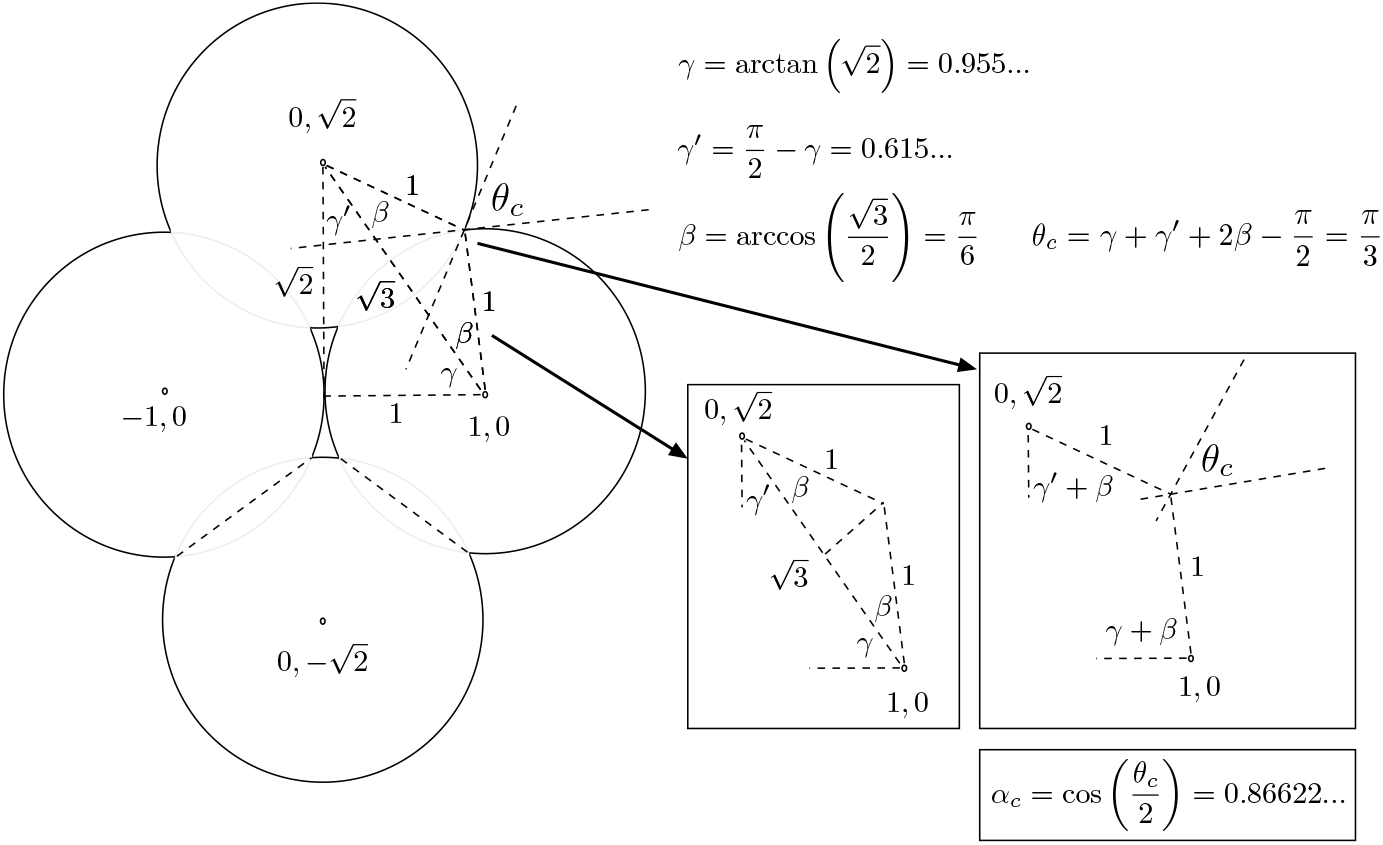
Computing the angle and the *α* triggering the internal contact and, therefore, rigidity.

## III. ADHESION-INDUCED COLLAPSE OF POROSITY

In this section we will show how the rigidification for values *α* < *α*_*c*_ explained above also entails the emergence/collapse of porosity of, that is, that all the interstitial voids become closed in the minimum energy state.

### A. Porosity

To study the effect of the reduction of the *α* on the porosity of the tissue, we follow a strategy based again on the fact that, in the 2D projection, cells in contact can be treated as two disks whose contact region is the bisection of intersection between both. We consider a starting point where two disks of radius R are separated by a distance 2R, having, therefore, only one contact point –see Suppl. Note Fig. (4). One can study the evolution of *α* by applying the following rescaling:

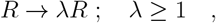

while keeping the distance of the centers of mass invariant. We refer to λ as the *scaling parameter*. In that context, the actual units become rescaled. However, there is no need to unfold the scaling, as long as we are interested in i) Cell fraction and ii) Intersection angles to infer the critical *α*’s derived from the Young-Dupré relation. Metaphorically speaking, we are inflating the cells while keeping the centers of mass static. Considering an arbitrary radius, for λ ≥ 1, we have that the intersection angle between two adjacent cells can be inferred as:

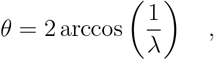

from which we conclude that:

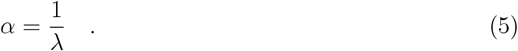

**Suppl. Note Fig. 4:**
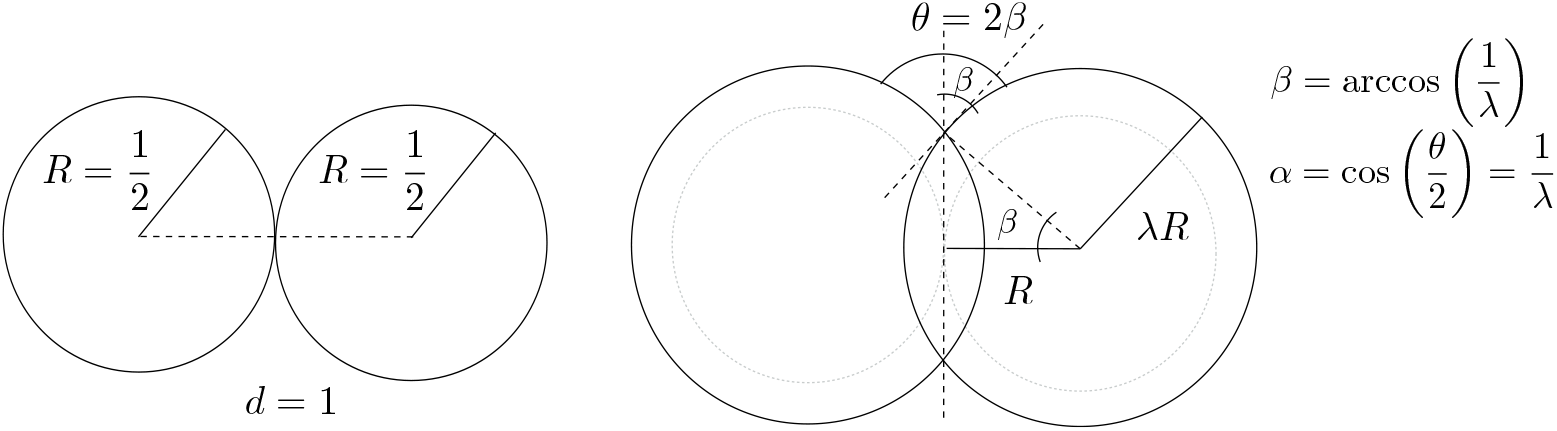
Schema detailing the use of the scaling parameter *λ*.

In Suppl. Note Fig. (4) we detail this observation.

The key point in the computations is to find at which point a 3-cell junction is formed –see Suppl. Note Fig. (5). Indeed, when 3-cell junctions are favoured, cells will tend to pack in confluent regions. In particular, using the scaling parameter λ, one can see that:

For 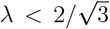, 3 cells in contact forming a triangle do not create spontaneously a 3-cell junction

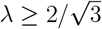, 3 cells in contact forming a triangle create spontaneously a 3-cell junction

**Suppl. Note Fig. 5:**
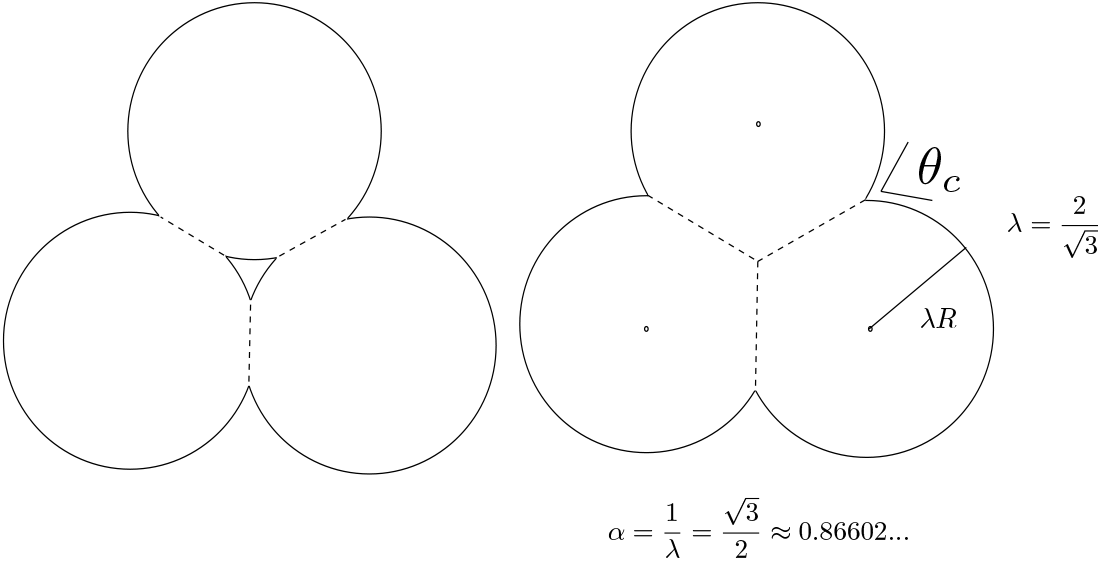
Collapse of the interstitial network: Computing the critical *α* for closing a three cell junction hole using the rescaling strategy described in the theorem.

To see that, we reason as follows: As long as the radius of the cell is larger than the distance from any of the cells to the centroid of the triangle, the gap is closed. In consequence, to compute the critical λ beyond which the hole is closed, the only thing we need to know is the distance between any cell of to the centroid of the triangle. Considering that the 3-cell arrangement is symmetrically distributed, this critical radius (R_*c*_) turns out to be:

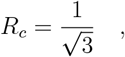

which implies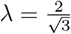. In Suppl. Note Fig.(5) we show the reasoning schematically. Thanks to equation (5) we can map this rescaled radius into the physically significant parameter *α*, leading to a critical point for confluency *α*_*c*_ of:

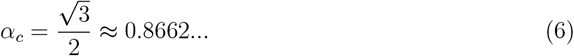

which, interestingly, coincides with the critical point of rigidity. This implies that *the emergence of adhesion-induced rigidity, as computed in section IIB of this supp. note, occurs at the same critical point of adhesion strength as the emergence of the adhesion-induced confluency/collapse of porosity*. Simulations using large-scale disordered arrangements of cells simulating tissues show a behaviour consistent with these predictions. Equivalent results for the threshold for confluency predicted in eq. (6) were previously reported in [8, 9]. We here extended the scope of the result using numerical simulations with arbitrary cell tilings and showing that the phenomenon of pore closing –and, in consequence, the transition to confluency– is expected in generic cell arrays. In addition, the derivation strategy allows us to compute the maximum cell fractions for all values of *α*, which is the target of the next subsection.

**Suppl. Note Fig. 6:**
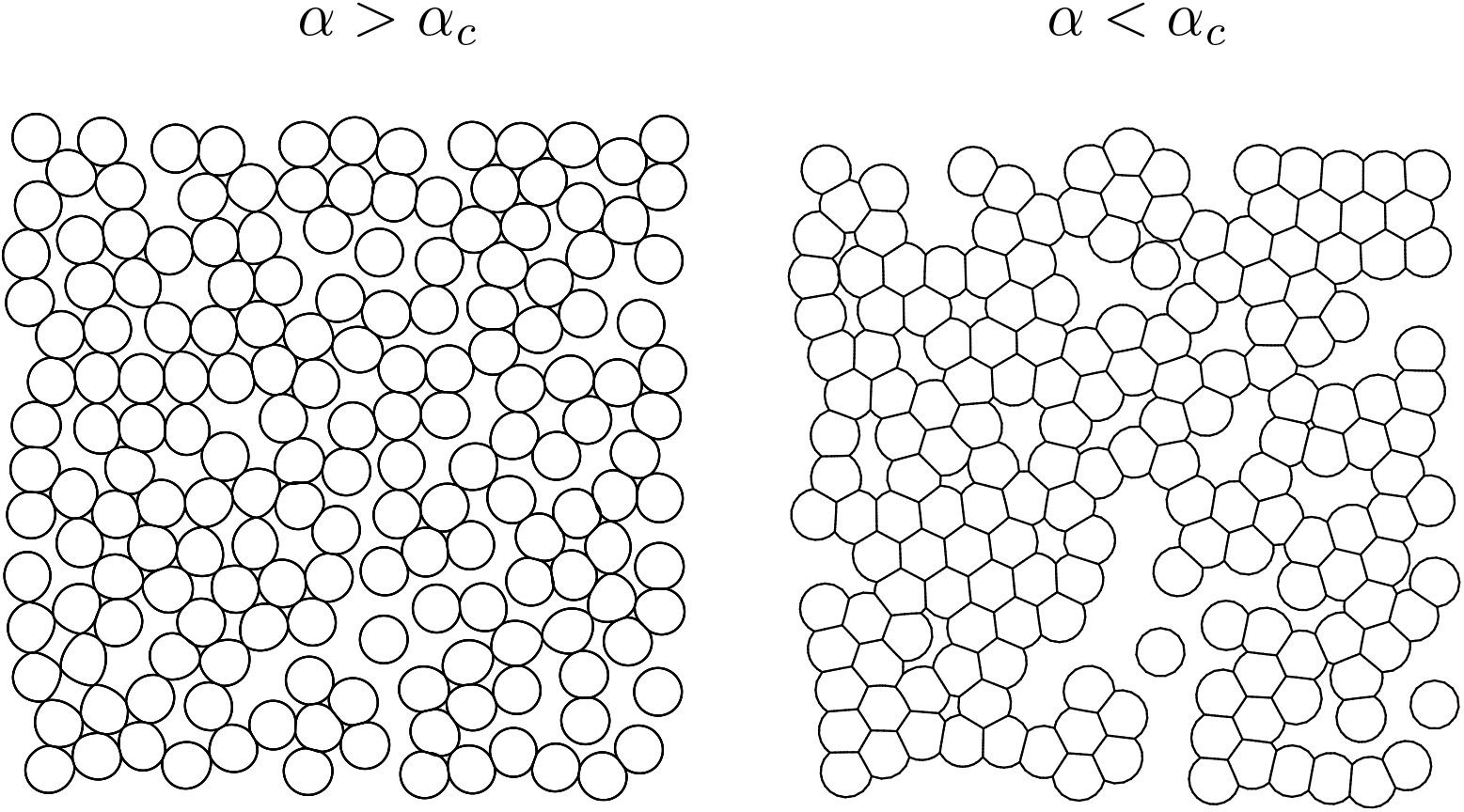
Simulations of real tissues at *α > α*_*c*_ and *α < α*_*c*_. We observe that, below *α*_*c*_ (right), almost all 3-cell junctions are closed. We used the same cell underlying arrangement for both simulations. *α*’s used: *α* = 0.93 (left), *α* = 0.83 (right).

### B. Maximal cell fractions

Start with a hexagonal tiling of hard disks of equal radius, as it is known that this is the arrangement leading to the maximal disk density. We then apply the following strategy: For each triangular junction, we consider the surface of triangle formed by the three centers of mass of three cells and how this is getting filled along the increase of λ. Considering the fact that the triangle contains 1/6 of each of the 3 cells and the 1/2 of the intersection surface shared by each pair of cells, we have that the relative surface, *ϕ*, occupied by the cells within the triangle is:

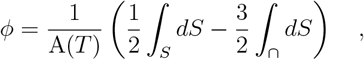

We assume that the circles for *α* = 1 have radius R = 1. This leads to an equilateral triangle of side 2, being its overall surface of the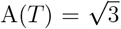. After applying a rescaling operation

R → λR, the other terms read:

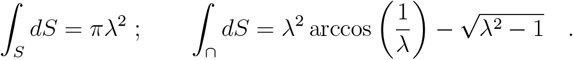

In terms of *α*, using that 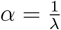

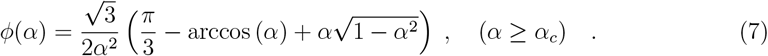

It is easy to check that *ϕ*(1) ≈ 0.907, as it is well known for the maximum density of the packing of hard disks of equal radius; and that *ϕ*(*α*_*c*_) = 1. The above equation tells us that full confluency is only possible, for tissues under force balance conditions, at values *α* ≤ *α*_*c*_. In Suppl. Note Fig. (7) the evolution of this maximum density is shown.

**Suppl. Note Fig. 7:**
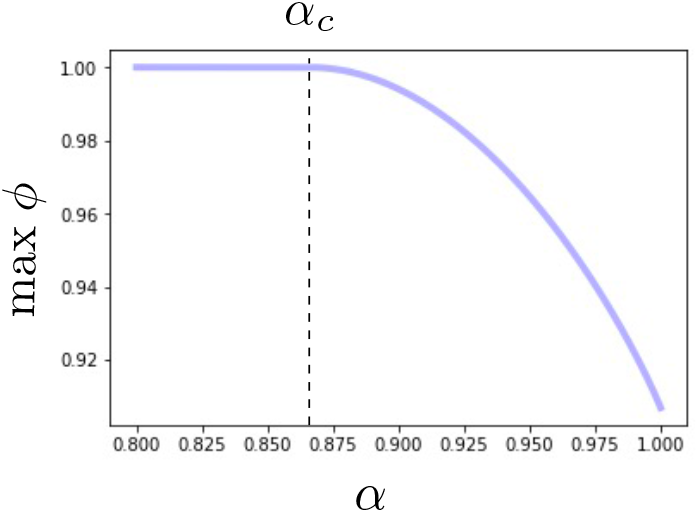
Maximum achievable cell fraction *ϕ* as a function of *α* according to eq. (7).

## IV. NUMERICAL SIMULATIONS

Numerical simulations have been performed using the C based software Surface Evolver version 2.70 [10]. To study the evolution of a 2D cell tiling using Surface Evolver, it is necessary to enter an initial *mosaic* –eventually with non-covered regions– specifying the location of the cells, a primary polygon-like geometry and the initial topology of cell-cell contacts. Later this initial condition will evolve 1/ By increasing the resolution of the perimeter of the cells –thereby achieving realistic geometries– and 2/ Optimizing the global geometry of individual cells and size of cell-cell contacts according to a global soap-bubble-like Hamiltonian like the one described in equation (3), in a relaxation process towards the desired *α* parameter.

Throughout this research we do not consider the action of external pulling - or pushing forces and, thereby, we expect the tilings to be in equilibrium with respect the soap-bubble Hamiltonian.

### A. Random seeds for arbitrary cell tilings

The definition of the initial random seed that will be further optimized to the desired *α* can be performed following different algorithmic procedures: 1/ The Lubachevsky-Stillinger algorithm, indicated to study jamming of hard disks 2/ Random location of disks in a plane, indicated to generate floppy –subcritical– cell arrangements with already existing contacts and 3/ a network-like based algorithm, in which the network topology is a-priori given, indicated for fine-grained exploration towards confluency. The density of these initial seeds is initially parametrized through an estimation. Since the optimization process may induce variations, the cell density is re-computed at the end of the process.

#### 1. Lubachevsky-Stillinger algorithm (LSA) –Jamming

The Lubachevsky-Stillinger (compression) algorithm (LSA) is the standard numerical method used to simulate the physical process of compressing an assembly of hard disks that eventually leads to hard-sphere-like jamming [11]. In this research we used the C++ based implementation provided in [12]. Although LSA method allows us to generate random tilings at different densities, most of the contacts between cells are formed at the limit of the jamming transition *ϕ*_*c*_ ≈ 0.84. In consequence, the LSA may not be indicated for producing the seeds for random tilings at subcritical densities.

#### 2. Random disk spreading (RDS) –Topological consequences of changes in *α*

The method described below allows us to generate random tilings with subcritical target densities *ϕ*_*T*_, and is based on the RSA algorithm reported in [13] to generate subcritical random disk arrays. This method is indicated for the study of rigidification of previously floppy structures upon decrease of *α*. The steps followed in the algorithm are described below:

- Over an a-priori defined square of length L –in units of cell diameter D = 2R, where R is the average cell radius– send N^′^ random possible coordinates 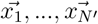.
- The sequential process of generation of random coordinates is subject to a selection criteria: If when generating the k-th random coordinate an already existing coordinate 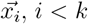 is such that 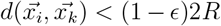 this coordinate 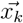 is discarded, as it would lead to a large overlapping pair of disks.
- If the number of accepted generated coordinates N (N ≤ N^′^) reaches a value such that, *ϕ*_*T*_ ≤ N*ϕ*R^2^/L^2^, where *ϕ*_*T*_ is the target density, the process stops, since the target density has been achieved.
- If after a long number of iterations, the target density cannot be achieved, we perform a random search along the area identifying possible empty spaces that can be filled using the previous distance conditions. Note that for densities close to *ϕ*_*c*_ this process is harder and harder and becomes technically unfeasible for *ϕ* ≈ *ϕ*_*c*_, where the LSA or the network based algorithm for generating tilings defined below are more indicated.
- We build a collection C_1_, …C_*N*_ of disks centered on each of the accepted coordinates 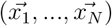. For any pair of disks C_*i*_, C_*k*_ such that 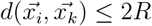 we define a contact.

Since the presence of overlaps and further changes on adhesion may slightly alter the cell fraction, we may encounter the situation by which the actual achieved density *ϕ* < *ϕ*_*T*_. If this happens, we need to generate more disks, refining the halting condition and rewriting it as *ϕ* ≤ (N + δ)*ϕ*R^2^/L^2^ to achieve the desired densities *ϕ* ≈ *ϕ*_*T*_ after optimization. The algorithm can be extended for polydisperse tilings.

#### 3. Network-based tilling generation (NTG) –transition to confluency

The following method is indicated for the generation of dense tilings and/or when we want to study the transition from non-confluent to confluent tissues upon decrease of *α*. The steps are the following:

- Generate a regular triangular lattice with a given size –side L cells with radius R. The position of the N nodes is 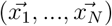. In this setting, adjacent nodes are at exactly distance 2R.
- Remove some sites with probability p, computed from the desired target density *ϕ*_*T*_
- Introduce noise in the geolocalization of the nodes. Position 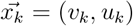 is replaced by 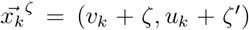 where ζ, ζ^′^ are random numbers drawn from a gaussian distribution centered at 0 whose standard deviation is σ = ϵR.
- For each 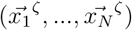 draw a circle of radius R around it.
- For each pair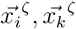, establish a contact if 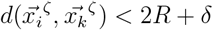.

### B. Adhesion-induced rigidification

We detail here how initially floppy tilings at subcritical target density *ϕ*_*T*_ can become generically rigid by decreasing the parameter *α* using the Surface Evolver software (version 2.70) [10]. Since we are working with subcritical densities and we want to observe changes in the topology, we use the above described RDS method to generate the seed which will define the starting point of the optimization process.

The parameter t sets the minimum segment length forming a polygon –which represents a cell–, thus controlling also the number of segments a cell is made of. As in previous sections, we consider 2D projections of cell arrangements, where each cell is described by a set of vertices. The method followed in this case is:

- Generate a random, subcritical tiling with target density *ϕ*_*T*_ < *ϕ*_*c*_ and the desired size with the RDS method.
- Stabilize the tiling in the hard-sphere regime (*α* = 1).
- Decrease *α* in a quasi-static way until the desired *α* is reached. In this step, the mechanism of re-meshing implemented by the Surface Evolver acts by alternatively reducing and increasing the resolution of the geometry of the cells. This is done by merging all segments lower than a certain length (t parameter) and subsequently dividing such segments by 2. With this, the optimization algorithm can explore the potential configurations minimizing the global surface energy in an efficient way. According to our simulations, working with very fine grained geometries (low t parameters) all the time may trap the system in local optima. In consequence, we need to introduce sporadically large merging events (large t’s) to trigger topological changes. To preserve the high resolution of the simulation but, at the same time, allow topological rearrangements, we apply the re-meshing t-parameter as following a Weibull function:

**Suppl. Note Fig. 8:**
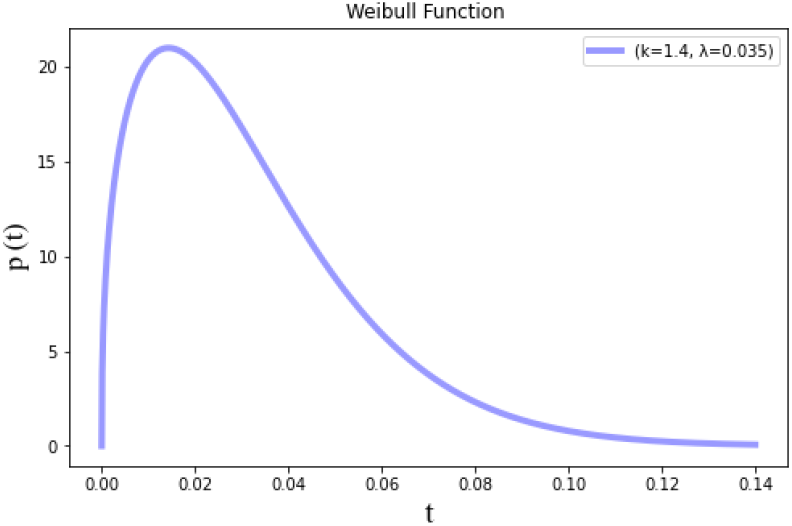
Weibull function for *k*=1.4 and *λ*=0.035.

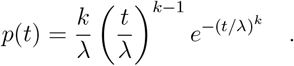

This allows us to perform most of the re-meshing events around a well defined mean, keeping high resolution, but, from time to time, massive re-meshing events allow topo-logical rearrangements.

When the targeted *α* is reached and no changes on the energy are appreciated throughout successive rounds of optimization, we end the process. The result is a tiling with the desired *α* and approximately the target density *ϕ*_*T*_. Since the density can be altered, it is recomputed after the optimization process.

### C. Adhesion-induced confluency

To test the mathematical prediction on the emergence of adhesion-induced confluency in tissues at *α*_*c*_ ≈ 0.866 (see the mathematical section), we perform several simulations considering different cell arrangements. As a starting point, we study one of the simplest topologies: a triangular motif of 3 symmetrically distributed cells of equal radius in contact. The simulation starts with cells modeled as non-adhesive disks at *α* = 1, which is progressively decreased to *α* = 0.7. Using the same initial configuration, we test different values of the resolution parameter t. Since the computation of the angle formed requires high resolution –to be as close as possible to a circle– we need to use very low t values. For these values of t we observe that at *α*_*c*_ the interstitial hole between cells is closed, leading to the formation of a TCC in which the three cells involved share a vertex.

To extend the analysis to tissues, we generate both ordered and disordered arrays of cells (of approximately 10×10 cells in size). We use the NTG method, described in the previous section, to construct the seed for the simulations. Simulations of the evolution of these tilings upon reduction of *α* are performed for different t analogously to the simple triangular case (from *α* = 1 up to *α* = 0.7). The percentage of tricellular contacts in an cell tiling is then:

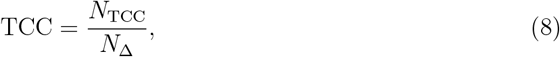

where N_TCC_ is the number of closed interstitial holes between 3 cells and N_Δ_ is the number of triangles in the contact network, respectively, which can be computed directly from the adjacency matrix of the network A, defined as:

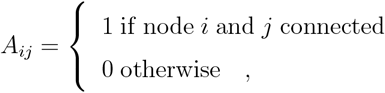

1 if node i and j connected 0 otherwise,

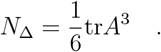

In both arrangements, for t = 0.01 a sharp transition occurs in the percentage of TCCs in the proximity of *α*_*c*_, indicating that below *α*_*c*_ the majority of holes are closed, in agreement with the mathematical predictions-For t < 0.01 this transition is not observed, possibly due to the existence of local minima trapping the system due to the small size of the segments.

### D. Computing the rigid clusters

Identification of floppy and rigid areas of the cell-cell contact networks was performed using pebble.py [15], available at https://github.com/coldlaugh/pebblegame-algorithm/blob/master/pebble.pyx.

## Notes

### Competing Interest Statement

The authors have declared no competing interest.

